# Genomic Prediction of Complex Disease Risk

**DOI:** 10.1101/506600

**Authors:** Louis Lello, Timothy G. Raben, Soke Yuen Yong, Laurent CAM Tellier, Stephen D.H. Hsu

## Abstract

We construct risk predictors using polygenic scores (PGS) computed from common Single Nucleotide Polymorphisms (SNPs) for a number of complex disease conditions, using L1-penalized regression (also known as LASSO) on case-control data from UK Biobank. Among the disease conditions studied are Hypothyroidism, (Resistant) Hypertension, Type 1 and 2 Diabetes, Breast Cancer, Prostate Cancer, Testicular Cancer, Gallstones, Glaucoma, Gout, Atrial Fibrillation, High Cholesterol, Asthma, Basal Cell Carcinoma, Malignant Melanoma, and Heart Attack. We obtain values for the area under the receiver operating characteristic curves (AUC) in the range ~ 0.58 – 0.71 using SNP data alone. Substantially higher predictor AUCs are obtained when incorporating additional variables such as age and sex. Some SNP predictors alone are sufficient to identify outliers (e.g., in the 99th percentile of PGS) with 3 – 8 times higher risk than typical individuals. We validate predictors out-of-sample using the eMERGE dataset, and also with different ancestry subgroups within the UK Biobank population. Our results indicate that substantial improvements in predictive power are attainable using training sets with larger case populations. We anticipate rapid improvement in genomic prediction as more case-control data become available for analysis.

## 1 Introduction

Many important disease conditions are known to be significantly heritable [1]. This means that genomic predictors and risk estimates for a large number of diseases can be constructed if enough case-control data is available. In this paper we apply L1-penalized regression (LASSO) to case-control data from UK Biobank [2] (UKBB) and construct disease risk predictors. Similar techniques have been used for phenotype prediction in plant and animal genomics, as described below, but are less familiar in the context of human complex traits and disease risks.^1^ In earlier work [9], we applied these methods to quantitative traits such as height, bone density, and educational attainment. Our height predictor captures almost all of the expected heritability for height and has a prediction error of roughly a few centimeters. Similar methods have also been employed in previous work on case-control datasets [10, 11]

The standard procedure for evaluating the performance of a genomic predictor is to construct the receiver operating characterstic (ROC) curve and compute the area under the ROC curve (AUC) [12]. Recently, Khera et al. [13] constructed risk predictors for Atrial Fibrillation, Type 2 Diabetes, Breast Cancer, Inflammatory Bowel Disease, and Coronary Artery Disease (CAD). For these conditions, they obtained AUCs of 0.77, 0.72, 0.68, 0.63 and 0.81 respectively. Note, though, that additional variables such as age and sex are used to obtain these results. When common SNPs alone are used in the predictors, the corresponding AUCs are smaller. For example, [14] obtain an AUC of 0.64 for CAD using SNPs alone – compared with 0.81 with inclusion of age and sex. (See also [15] for a CAD meta-analysis that also predicts risk stratification.)

Among the disease conditions studied here are Hypothyroidism, Hypertension, Type 1 and 2 Diabetes, Breast Cancer, Prostate Cancer, Testicular Cancer, Gallstones, Glaucoma, Gout, Atrial Fibrillation, High Cholesterol, Asthma, Basal Cell Carcinoma, Malignant Melanoma and Heart Attack. We obtain AUCs in the range 0.580 – 0.707 (see Table 1), using SNP data alone. Substantially higher AUCs are obtained by incorporating additional variables such as age and sex. Some SNP predictors alone are sufficient to identify outliers (e.g., in the 99th percentile of PGS) with, e.g., 3 – 8 times higher risk than typical individuals. We validate predictors out-of-sample using the eMERGE dataset[16] (taken from the US population), and also with different ancestry subgroups within the UK Biobank population as done in [17].

**Table 1.**
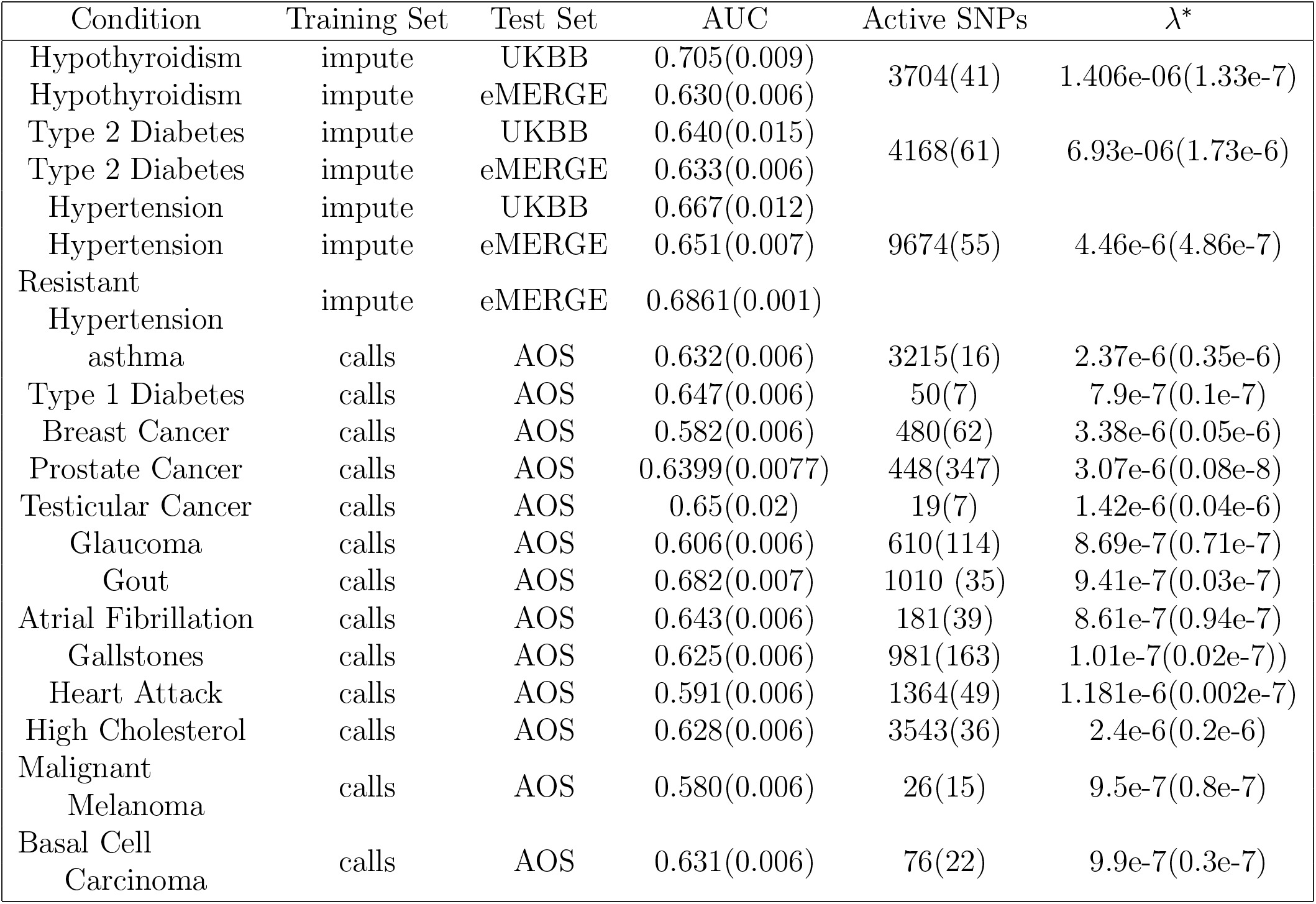
Table of genetic AUCs using SNPs only – no age or sex. Training and validating is done using UKBB data from either direct calls or imputed data to match eMERGE. Testing is done with UKBB, eMERGE, or AOS as described in Secs. 2 and Appendix D. Numbers in parenthesis are the larger of either a standard deviation from central value or numerical precision as described in Sec. 2. λ^*^ refers to the lasso λ value used to compute AUC as described in Sec. 2.

| Condition | Training Set | Test Set | AUC | Active SNPs | $\lambda^*$ |
| --- | --- | --- | --- | --- | --- |
| Hypothyroidism | impute | UKBB | 0.705(0.009) | 3704(41) | 1.406e-06(1.33e-7) |
| Hypothyroidism | impute | eMERGE | 0.630(0.006) |  |  |
| Type 2 Diabetes | impute | UKBB | 0.640(0.015) | 4168(61) | 6.93e-06(1.73e-6) |
| Type 2 Diabetes | impute | eMERGE | 0.633(0.006) |  |  |
| Hypertension | impute | UKBB | 0.667(0.012) | 9674(55) | 4.46e-6(4.86e-7) |
| Hypertension | impute | eMERGE | 0.651(0.007) |  |  |
| Resistant Hypertension | impute | eMERGE | 0.6861(0.001) |  |  |
| asthma | calls | AOS | 0.632(0.006) | 3215(16) | 2.37e-6(0.35e-6) |
| Type 1 Diabetes | calls | AOS | 0.647(0.006) | 50(7) | 7.9e-7(0.1e-7) |
| Breast Cancer | calls | AOS | 0.582(0.006) | 480(62) | 3.38e-6(0.05e-6) |
| Prostate Cancer | calls | AOS | 0.6399(0.0077) | 448(347) | 3.07e-6(0.08e-8) |
| Testicular Cancer | calls | AOS | 0.65(0.02) | 19(7) | 1.42e-6(0.04e-6) |
| Glaucoma | calls | AOS | 0.606(0.006) | 610(114) | 8.69e-7(0.71e-7) |
| Gout | calls | AOS | 0.682(0.007) | 1010 (35) | 9.41e-7(0.03e-7) |
| Atrial Fibrillation | calls | AOS | 0.643(0.006) | 181(39) | 8.61e-7(0.94e-7) |
| Gallstones | calls | AOS | 0.625(0.006) | 981(163) | 1.01e-7(0.02e-7)) |
| Heart Attack | calls | AOS | 0.591(0.006) | 1364(49) | 1.181e-6(0.002e-7) |
| High Cholesterol | calls | AOS | 0.628(0.006) | 3543(36) | 2.4e-6(0.2e-6) |
| Malignant Melanoma | calls | AOS | 0.580(0.006) | 26(15) | 9.5e-7(0.8e-7) |
| Basal Cell Carcinoma | calls | AOS | 0.631(0.006) | 76(22) | 9.9e-7(0.3e-7) |

Our analysis indicates that substantial improvements in predictive power are attainable using training sets with larger case populations. We anticipate rapid improvement in genomic prediction as more case-control data become available for analysis.

It seems likely that genomic prediction of disease risk will, for a number of important disease conditions, soon be good enough to be applied broadly in a clinical setting [18–21]. Inexpensive genotyping (e.g., roughly $50 per sample for an array genotype which directly measures roughly a million SNPs, and allows imputation of millions more) can identify individuals who are outliers in risk score, and hence are candidates for additional diagnostic testing, close observation, or preventative intervention (e.g., behavior modification).

We note the successful application of similar methods in genomic prediction of plant and animal phenotypes. Earlier studies have shown some success on complex human disease risk using much smaller datasets and a variety of methods [22–24]. Early work in this direction can be found in, for example, [25] (which highlights the utility of what were then referred to as dense marker data sets), [3, 26, 27] (genome-wide allele significance from association studies in additive models), [28–30] (regression analysis), and [31] (accounting for linkage disequilibrium). For more recent reviews, and the current status of these approaches for plant and animal breeding, see [32–34].

## 2 Methods and Data

The main dataset we use for training is the 2018 release of the UKBB [35]^2^. We use only genetically British individuals (as defined by UKBB using principal component analysis described in [36]) for training of our predictors. For out of sample testing, we use eMERGE data (restricted to self-reported white Americans) as well as self-reported white but non-genetically British individuals in UKBB. The specific eMERGE data set used here refers to data obtained from dbGaP, under accession phs000360.v3.p1.(https://www.ncbi.nlm.nih.gov/projects/gap/cgi-bin/study.cgi?study_id=phs000360.v3.p1) We refer to the latter testing method as Ancestry Out of Sample (AOS) testing: the individuals used are part of the UKBB dataset, but have not been used in training and differ in ancestry from the training population.^3^

We construct linear models of genetic predisposition for a variety of disease conditions^4^. The phenotype data describes case-control status where cases are defined by whether the individual has been diagnosed for, or self-reports, the disease condition of interest. Our approach is built from previous work on compressed sensing [9, 44, 45]. In this earlier work we showed that matrices of human genomes are good “sensing matrices” in the terminology of compressed sensing. That is, the celebrated theorems resulting in performance guarantees and phase transition behavior of the *L*_1_ algorithms hold when human genome data are used [46–50]. Furthermore, *L*_1_ penalization efficiently captures essentially all the expected common SNP heritability for human height, one of the most complex but highly heritable human traits [9]. It is for these reasons that we focus specifically on *L*_1_ methods in this paper. We do not exclude the possibility that other methods (e.g. [51]) may work as well or better. *However, our primary motivation is the construction of potentially clinically useful predictors, not methodological comparison between different algorithms.*

We note that there are robust Bayesian Monte Carlo approaches that can account for a wide variety of model features like linkage disequilibrium and variable selection. However, it has been noted that (so far) for human complex traits, these methods have only produced a modest increase in predictive power at the cost of large computation times [52]. Our methods are not explicitly Bayesian; we estimate posterior uncertainties in our predictor via repeated cross-validation.

For each disease condition, we compute a set of additive effects 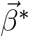 (each component is the effect size for a specific SNP) which minimizes the LASSO objective function:

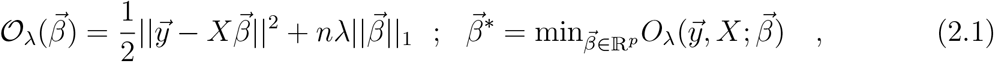

where *p* is the number of regressands, *n* is the number of samples, ||…|| means *L*_2_ norm (square root of sum of squares), ||…||_1_ is the *L*_1_ norm (sum of absolute values) and the term 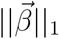 is a penalization which enforces sparsity of 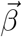. The optimization is performed over a space of 50,000 SNPs which are selected by rank ordering the p-values obtained from single-marker regression of the phenotype against the SNPs. The details of this are described in Appendix E.

Predictors are trained using a custom implementation of the LASSO algorithm which uses coordinate descent for a fixed value of λ. We typically use five non-overlapping sets of cases and controls held back from the training set for the purposes of in-sample cross-validation. For each value of λ, there is a particular predictor which is then applied to the cross-validation set, where the polygenic score is defined as (*i* labels the individual and *j* labels the SNP)

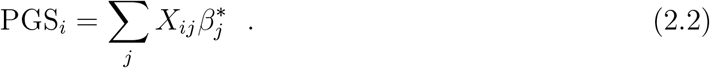

The term “polygenic score” typically refers to a simple measure built using results from single marker regression (e.g. GWAS), perhaps combined with p-value thresholding, and some method to account for linkage disequilibrium. Our use of penalized regression incorporates similar features – it favors sparse models (setting most effects to zero) in which the activated SNPs (those with non-zero effect sizes) are only weakly correlated to each other [9]. A thorough discussion of PGS construction is given in [53]. A brief overview of the use of single marker regression for phenotypes studied here is reviewed in Section D.

To generate a specific value of the penalization λ^*^ which defines our final predictor (for final evaluation on out-of-sample testing sets), we find the λ that maximizes AUC in each crossvalidation set, average them, then move one standard deviation in the direction of higher penalization (the penalization λ is progressively reduced in a LASSO regression). Moving one standard deviation in the direction of higher penalization errs on the side of parsimony^5^. These values of λ* are reported in Table 1, but further analysis shows that tuning λ to a value that maximizes the testing set AUC tends to match λ* within error. This is explained in more detail in Appendix E. The value of the phenotype variable *y* is simply 1 or 0 (for case or control status, respectively).

Scores can be turned into ROC curves by binning and counting cases and controls at various reference score values. The ROC curves are then numerically integrated to get AUC curves. We test the precision of this procedure by splitting ROC intervals into smaller and smaller bins. For several phenotypes this is compared to the rank-order (Mann-Whitney) exact AUC. The numerical integration, which was used to save computational time, gives AUC results accurate to ~ 1%^6^. For various AUC results the error is reported as the larger of either this precision uncertainty or the statistical error of repeated trials.

Finally we note that for the analysis of case-control phenotypes it is common to use logistic regression. We studied this approach for those of our phenotypes that also appear in [13], but found little to no difference in AUC or odds ratio results between linear and logistic regression. This might suggest that the data sets are highly constrained by the linear central region of the logistic function. Additionally, if we are simply interested in identifying genomes corresponding to *extreme* outliers, a linear regression can be more conservative.

## 3 Main Results

Figure 2 shows the evaluation of a predictor built using the LASSO algorithm. The LASSO outputs can be used to build ROC curves, as shown in Fig. 1, and in turn produce AUCs and Odds Ratios. Five non-overlapping sets of cases and controls are held back from the training set for the purposes of in-sample cross-validation. For each value of λ, there is a particular predictor which is then applied to the cross-validation set. The value of λ one standard deviation higher than the one which maximizes AUC on a cross-validation set is selected as the definition of the model. Models are additionally judged by comparing a non-parametric measure, Mann-Whitney data AUC, to a parametric prediction, Gaussian AUC.

Each training set builds a slightly different predictor. After each of the 5 predictors is applied to the in-sample cross-validation sets, each model is evaluated (by AUC) to select the value of λ which will be used on the testing set. For some phenotypes we have access to true out-of-sample data (i.e. eMERGE), while for other phenotypes we implement ancestry out-of-sample (AOS) testing using genetically dissimilar groups [17]. This is described in Appendices C and D. An example of this type of calculation is shown in Figure 2, where the AUC is plotted as a function of λ for Hypertension.

**Figure 1:**
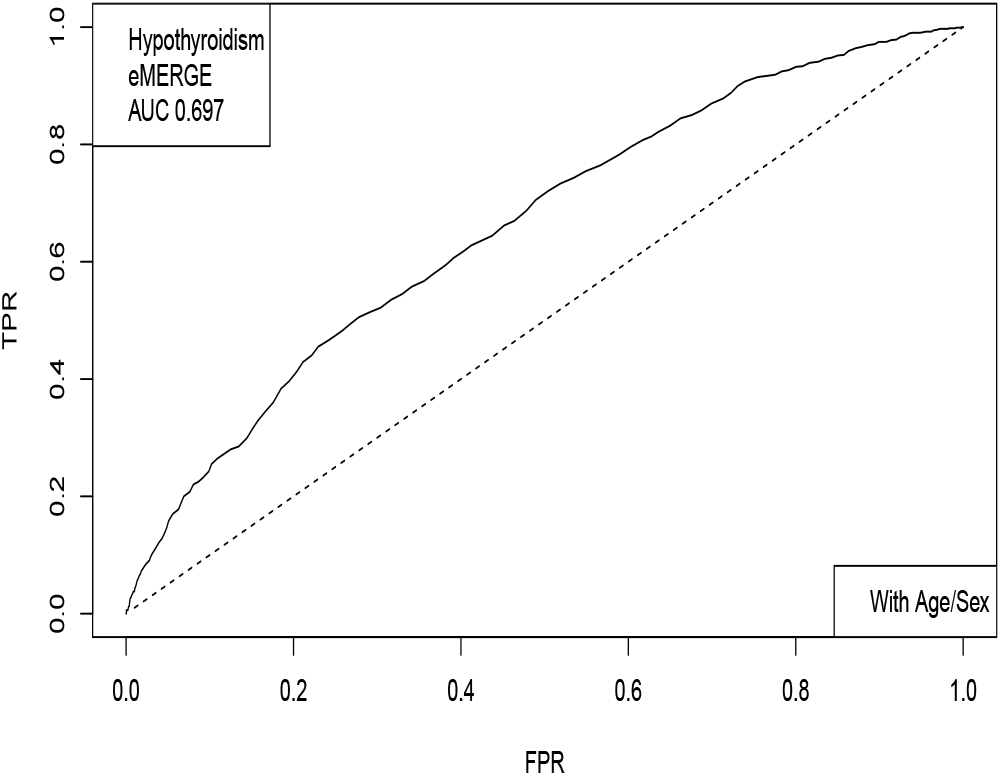
The receiver operator characteristic curve for case-control data on Hypothyroidism. This example includes sex and age as covariates.

**Figure 2:**
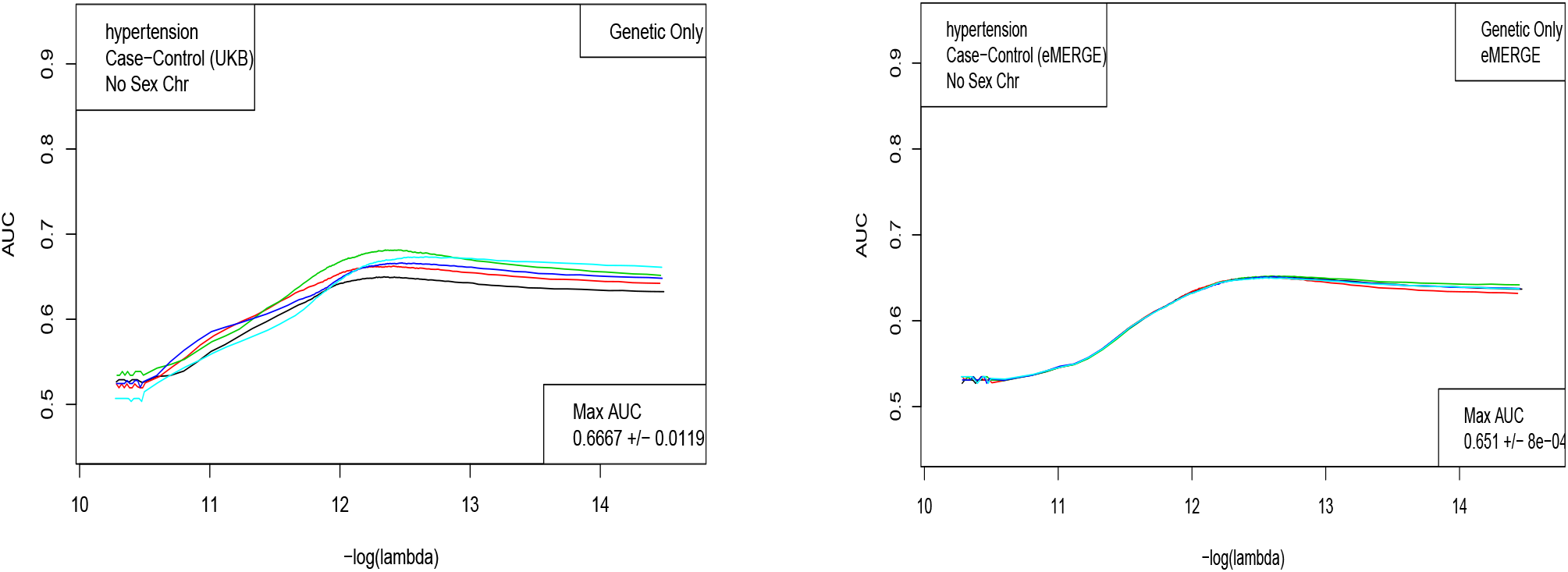
AUC computed on 5 holdback sets (1,000 each of cases and controls) for Hypertension, as a function of λ. A. UK Biobank and B. eMERGE.

Table 1 below presents the results of similar analyses for a variety of disease conditions. We list the best AUC for a given trait and the data set which was used to obtain that AUC.

In Figs. 3,4,5, and 6, the distributions of the polygenic score are shown for cases and controls drawn from the eMERGE dataset. In each figure, we show on the left the distributions obtained from performing LASSO on case-control data only, and on the right an improved polygenic score which includes effects from separately regressing on sex and age. The improved polygenic score is obtained as follows: regress the phenotype *y* = (1, 0) against sex and age, and then add the resulting model to the LASSO score. This procedure is reasonable since SNP state, sex, and age are independent degrees of freedom. In some cases, this procedure leads to vastly improved performance.

**Figure 3:**
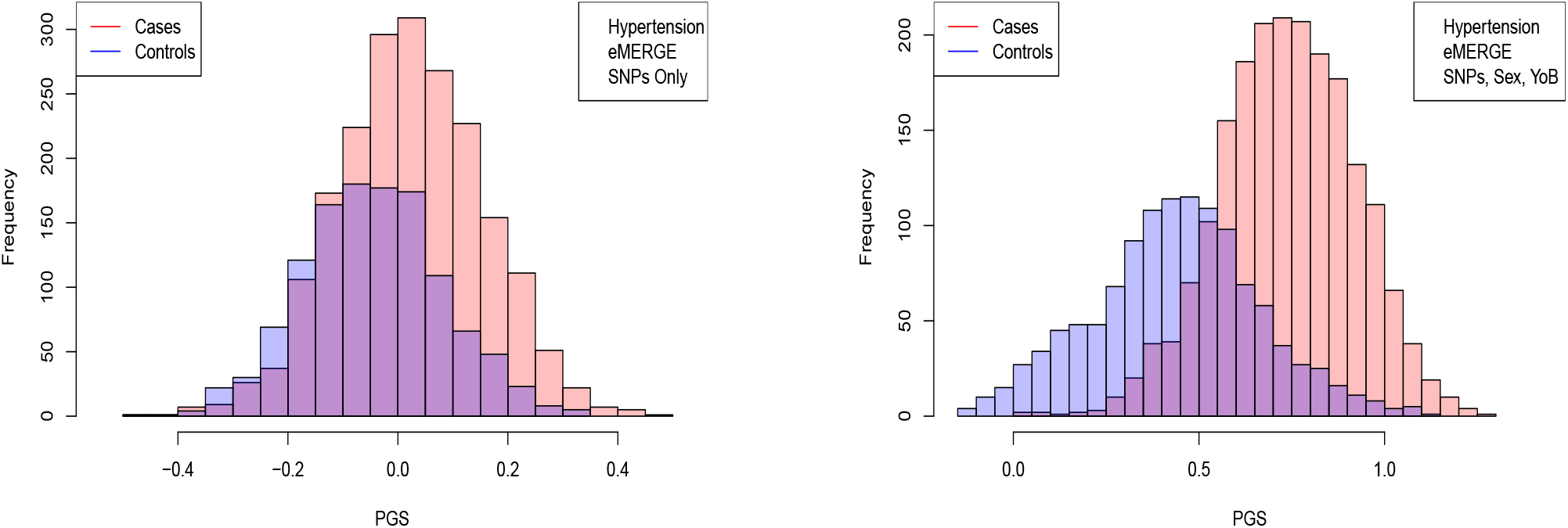
Distribution of PGS, cases and controls for Hypertension in the eMERGE dataset using SNPs alone and including sex and age as regressors.

**Figure 4:**
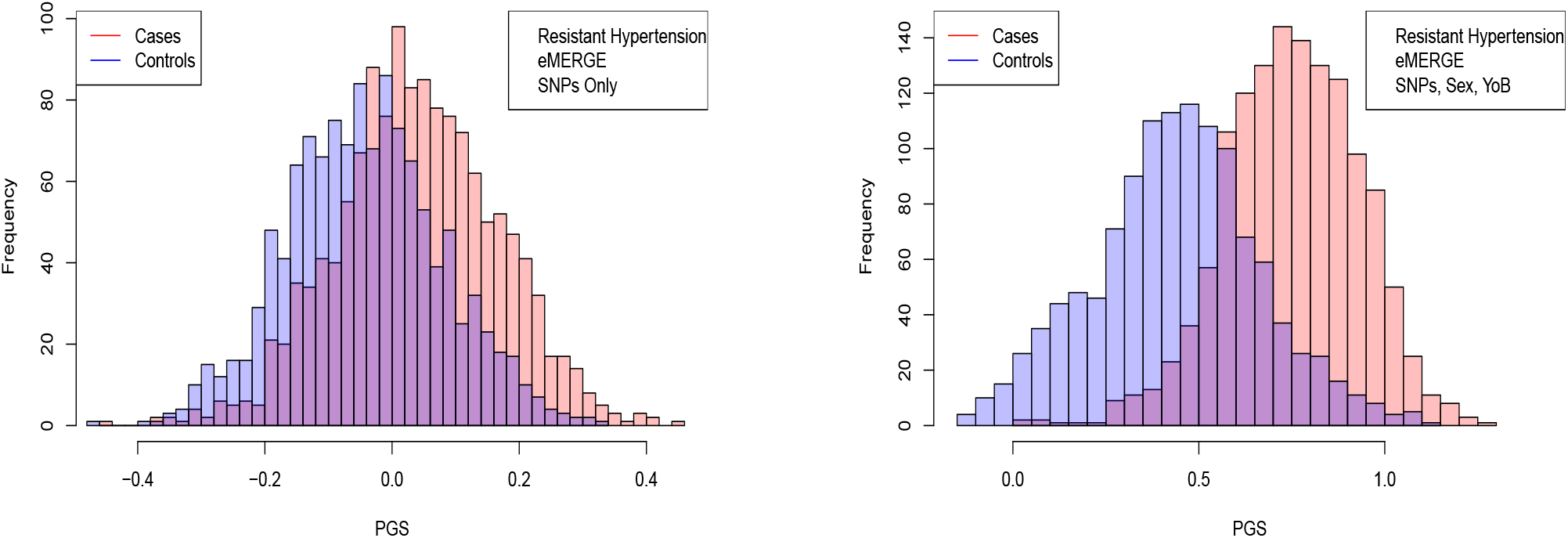
Distribution of PGS score, cases and controls for Resistant Hypertension in the eMERGE dataset using SNPs alone and including sex and age as regressors.

**Figure 5:**
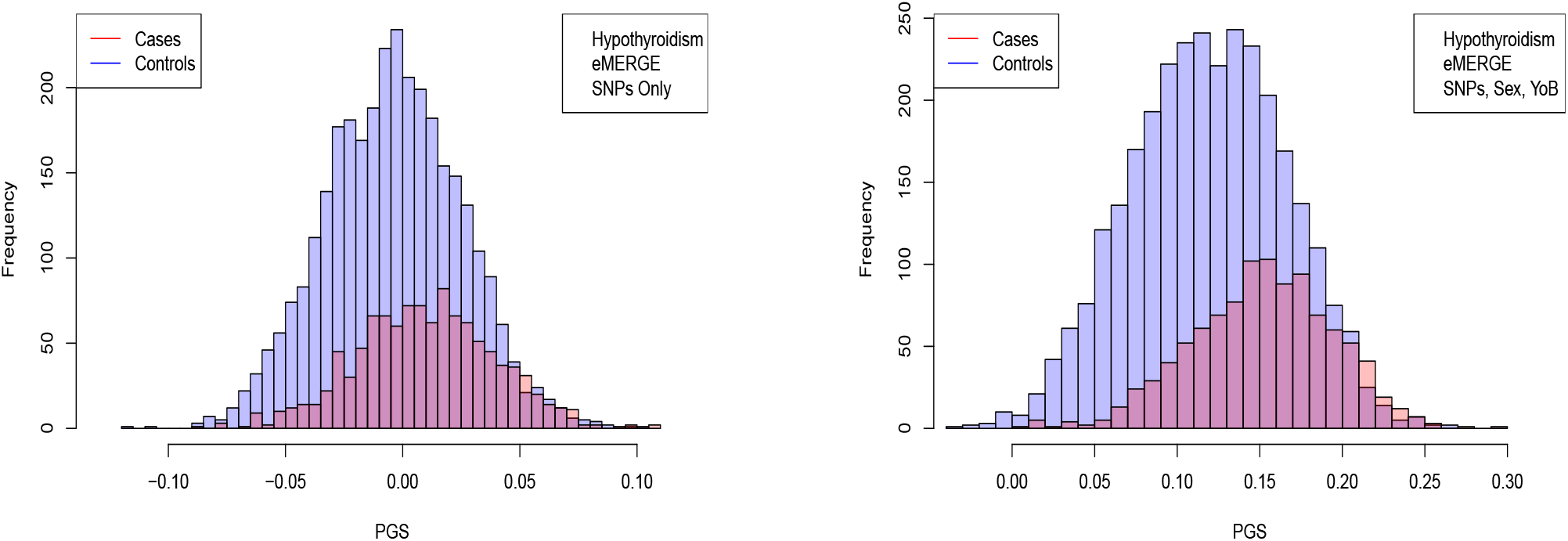
Distribution of PGS score, cases and controls for Hypothyroidism in the eMERGE dataset using SNPs alone and including sex and age as regressors.

**Figure 6:**
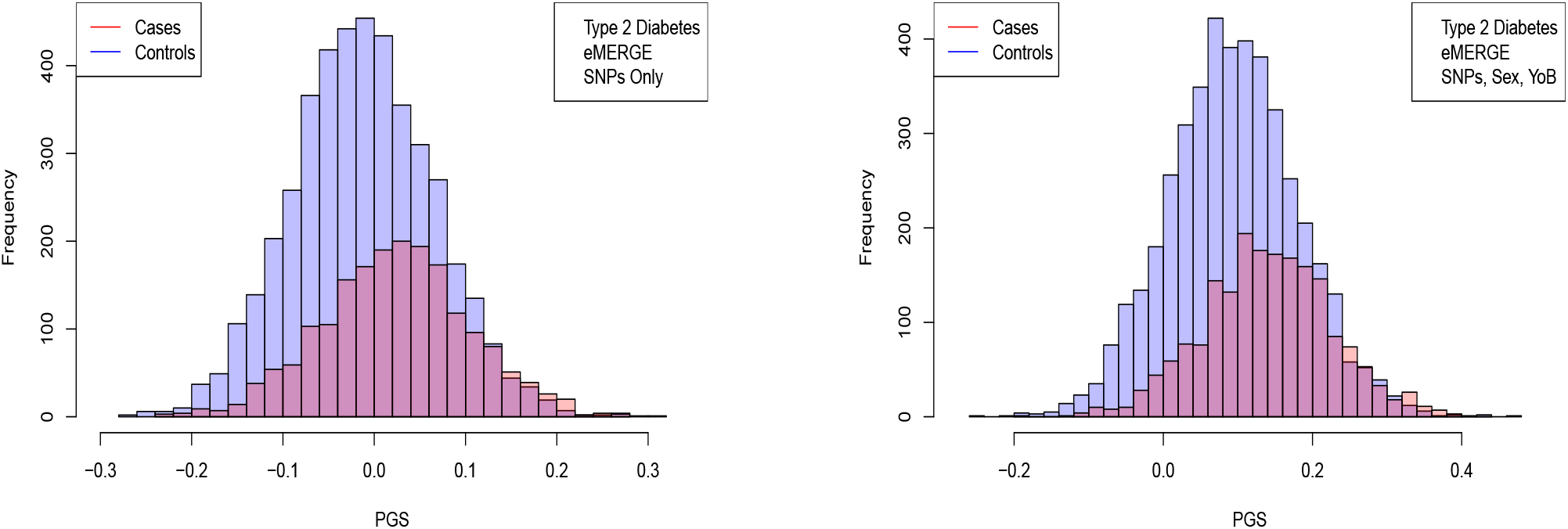
Distribution of PGS score, cases and controls for type 2 diabetes in the eMERGE dataset using SNPs alone and including sex and age as regressors.

The distribution of PGS among cases can be significantly displaced (e.g., shifted by a standard deviation or more) from that of controls when the AUC is high. At modest AUC, there is substantial overlap between the distributions, although the high-PGS population has a much higher concentration of cases than the rest of the population. Outlier individuals who are at high risk for the disease condition can therefore be identified by PGS score alone even at modest AUCs, for which the case and control normal distributions are displaced by, e.g., less than a standard deviation.

In table 2 we compare results from regressions on SNPs alone, sex and age alone, and all three combined. Performance for some traits is significantly enhanced by inclusion of sex and age information.

**Table 2:** AUCs obtained using sex and age alone, SNPs alone, and all three together.

| Condition | Test set | Age + Sex | Genetic Only | Age + Sex + genetic |
| --- | --- | --- | --- | --- |
| Hypertension | UKBB | 0.638 (0.018) | 0.667 (0.012) | 0.717 (0.007) |
| Hypothyroidism | UKBB | 0.695 (0.007) | 0.705 (0.009) | 0.783 (0.008) |
| Type 2 Diabetes | UKBB | 0.672 (0.009) | 0.640 (0.015) | 0.651 (0.013) |
| Hypertension | eMERGE | 0.818 (0.008) | 0.651 (0.007) | 0.851 (0.009) |
| Resistant Hypertension | eMERGE | 0.817 (0.008) | 0.686 (0.007) | 0.864 (0.009) |
| Hypothyroidism | eMERGE | 0.643 (0.006) | 0.630 (0.006) | 0.697 (0.007) |
| Type 2 Diabetes | eMERGE | 0.565 (0.006) | 0.633 (0.006) | 0.651 (0.007) |

For example, Hypertension is predicted very well by age + sex alone compared to SNPs alone whereas Type 2 Diabetes is predicted very well by SNPs alone compared to age + sex alone. In all cases, the combined model outperforms either individual model.

The results thus far have focused on predictions built on the autosomes alone (i.e. SNPs from the sex chromosomes are not included in the regression). However, given that some conditions are predominant in one sex over the other, it seems possible that there is a nontrivial effect coming from the sex chromosomes. For instance, 85% of Hypothyroidism cases in the UK Biobank are women. In table 3 we compare the results from including the sex chromosomes in the regression to using only the autosomes. The differences found in terms of AUC is negligible, suggesting that variation among common SNPs on the sex chromosomes does not have a large effect on Hypothyroidism risk. We found a similarly negligible change when including sex chromosomes for AOS testing.

**Table 3:**
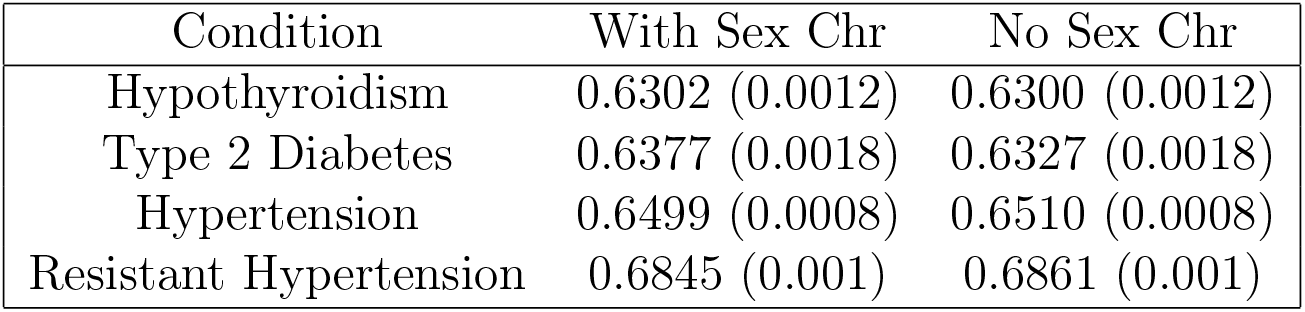
AUCs with and without SNPs from the sex chromosomes. All tested on eMERGE using SNPs as the only covariate.

Figs. 3,4,5, and 6 suggest that case and control populations can be approximated by two overlapping normal distributions. Under this assumption, one can relate AUC directly to the means and standard deviations of the case and control populations. If two normal distributions with means *μ*_1_,*μ*_0_ and standard deviations *σ*_1_,*σ*_0_ are assumed for cases and controls (*i* = 1, 0 respectively below), the AUC can be explicitly calculated via^7^

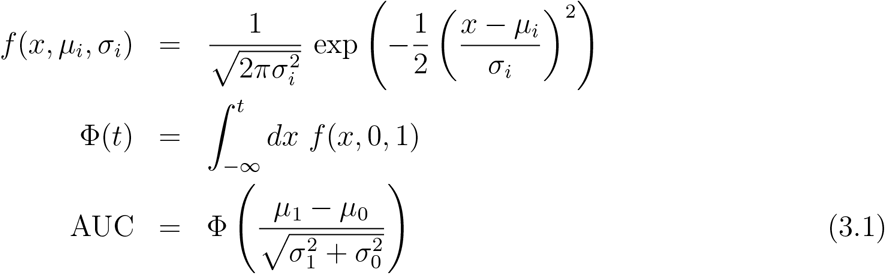

Under the assumption of overlapping normal distributions, we can compute the following odds ratio OR(*z*) as a function of PGS. OR(*z*) is defined as the ratio of cases to controls for individuals with PGS ≥ *z* to the overall ratio of cases to controls in the entire population. In the formula below, 1 = cases, 0 = controls.

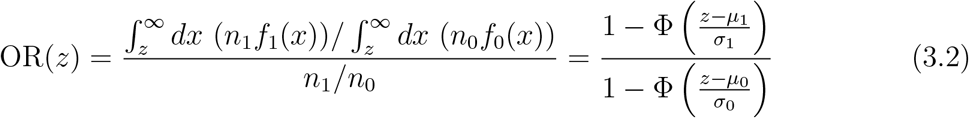

We compute means and standard deviations for cases and controls using the PGS distribution defined by the best predictor (by AUC) in the eMERGE dataset. We can then compare the AUC and OR predicted under the assumption of displaced normal distributions with the actual AUC and OR calculated directly from eMERGE data.

AUC results are shown in table 4, where we assemble the statistics for predictors trained on SNPs alone. In table 5 we do the same for predictors trained on SNPs, sex, and age.

**Table 4:**
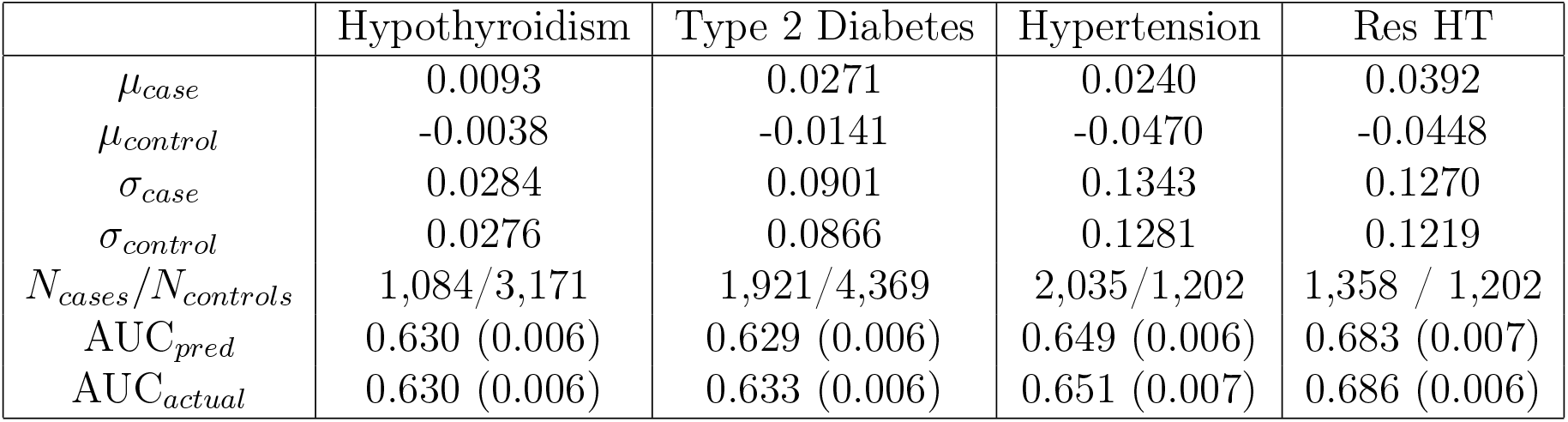
Mean and standard deviation for PGS distributions for cases and controls, using predictors built from SNPs only and trained on case-control status alone. Predicted AUC from assumption of displaced normal distributions and actual AUC are also given.

**Table 5:** Mean and standard deviation for PGS distributions of cases and controls, using predictors built from SNPs, sex, and age, and trained on case-control status alone. Predicted AUC from assumption of displaced normal distributions and actual AUC are also given.

|  | Hypothyroidism | Type 2 Diabetes | Hypertension | Res HT |
| --- | --- | --- | --- | --- |
| $\mu_{case}$ | 0.1516 | 0.1431 | 0.7377 | 0.7525 |
| $\mu_{control}$ | 0.1185 | 0.0924 | 0.4375 | 0.4366 |
| $\sigma_{case}$ | 0.0437 | 0.0948 | 0.1829 | 0.1830 |
| $\sigma_{control}$ | 0.0474 | 0.0943 | 0.2250 | 0.2258 |
| $N_{cases}/N_{controls}$ | 1,035/3,047 | 1,921/4,369 | 2,000/1,196 | 1,331/1,196 |
| $AUC_{pred}$ | 0.696 (0.007) | 0.648 (0.006) | 0.850 (0.009) | 0.862 (0.009) |
| $AUC_{actual}$ | 0.697 (0.007) | 0.651 (0.007) | 0.852 (0.009) | 0.864 (0.009) |

The results for odds ratios as a function of PGS percentile for several conditions are shown in figures 7,8,9,10. Note that each figure shows the results when 1) performing LASSO on case-control data only and 2) adding a regression model on sex + age to the LASSO result. The red line is what one obtains using the assumption of displaced normal distributions (i.e., Equation 3.2). Overall there is good agreement between directly calculated odds ratios and the red line. Odds ratio error bars come from 1) repeated calculations using different training sets and 2) by assuming that counts of cases and controls are Poisson distributed. (This increases the error bar or estimated uncertainty significantly when the number of cases in a specific PGS bin is small.)

**Figure 7:**
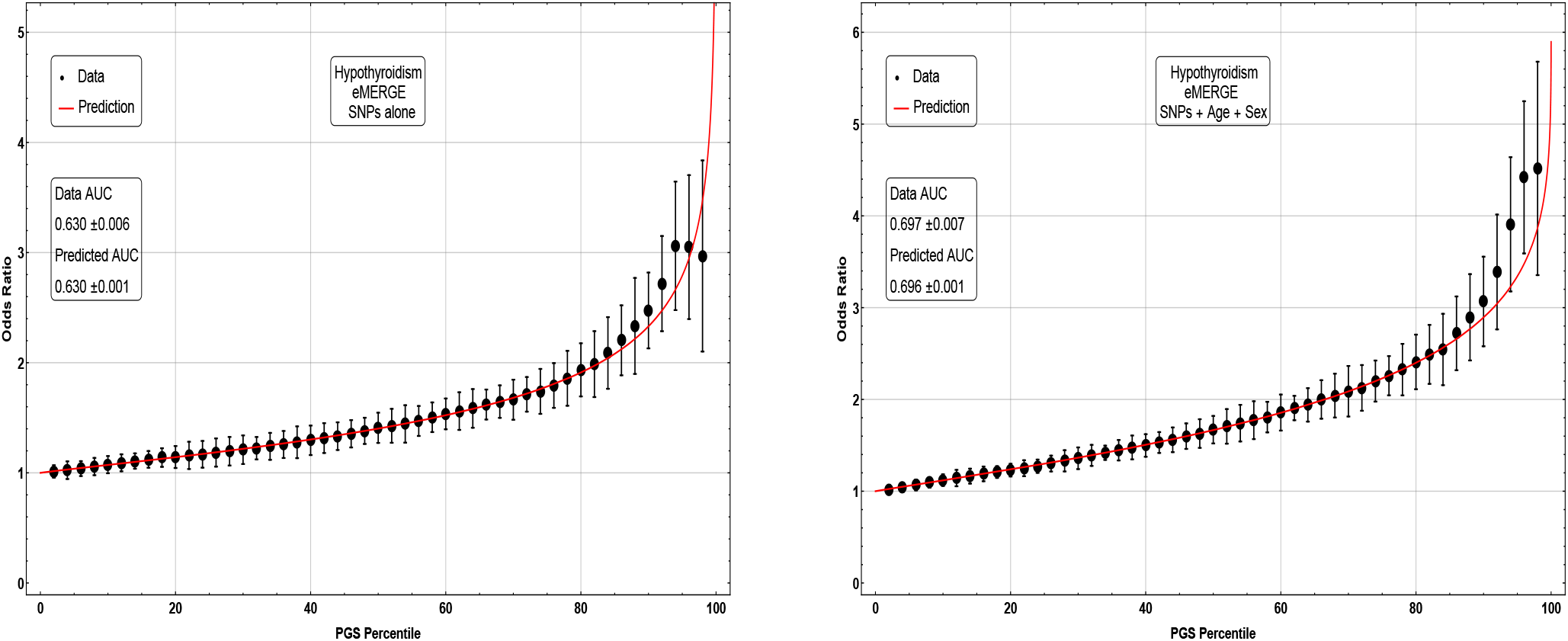
Odds ratio between upper percentile in PGS and total population prevalence in eMERGE for Hypothyroidism with and without using age and sex as covariates.

**Figure 8:**
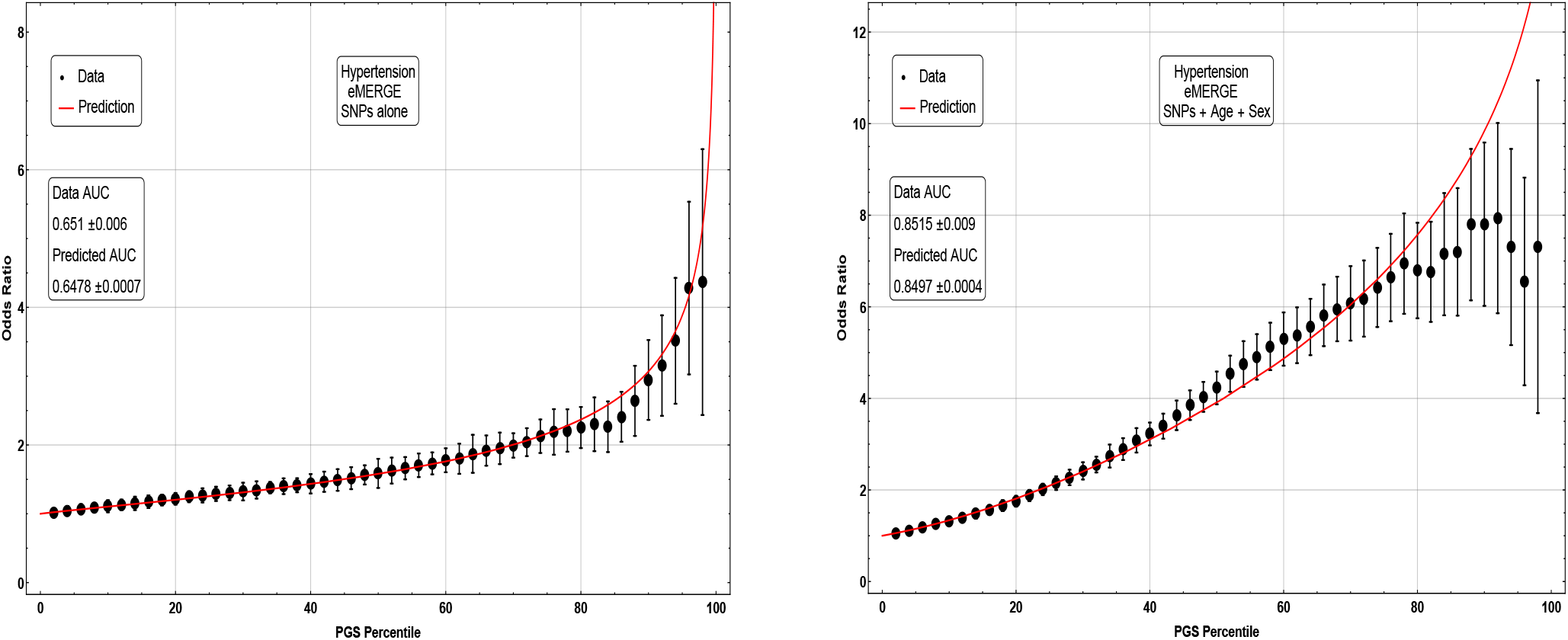
Odds ratio between upper percentile in PGS and total population prevalence in eMERGE for Hypertension with and without using age and sex as covariates.

**Figure 9:**
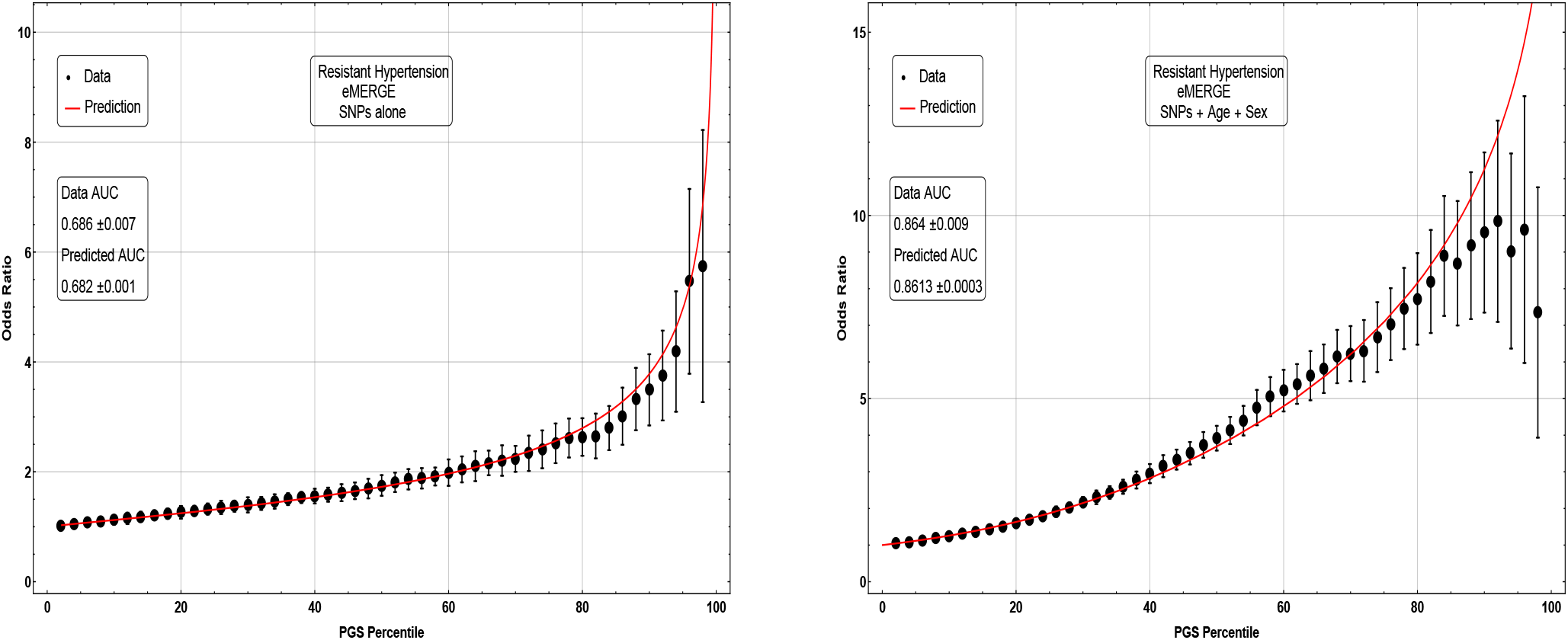
Odds ratio between upper percentile in PGS and total population prevalence in eMERGE for Resistant Hypertension with and without using age and sex as covariates.

**Figure 10:**
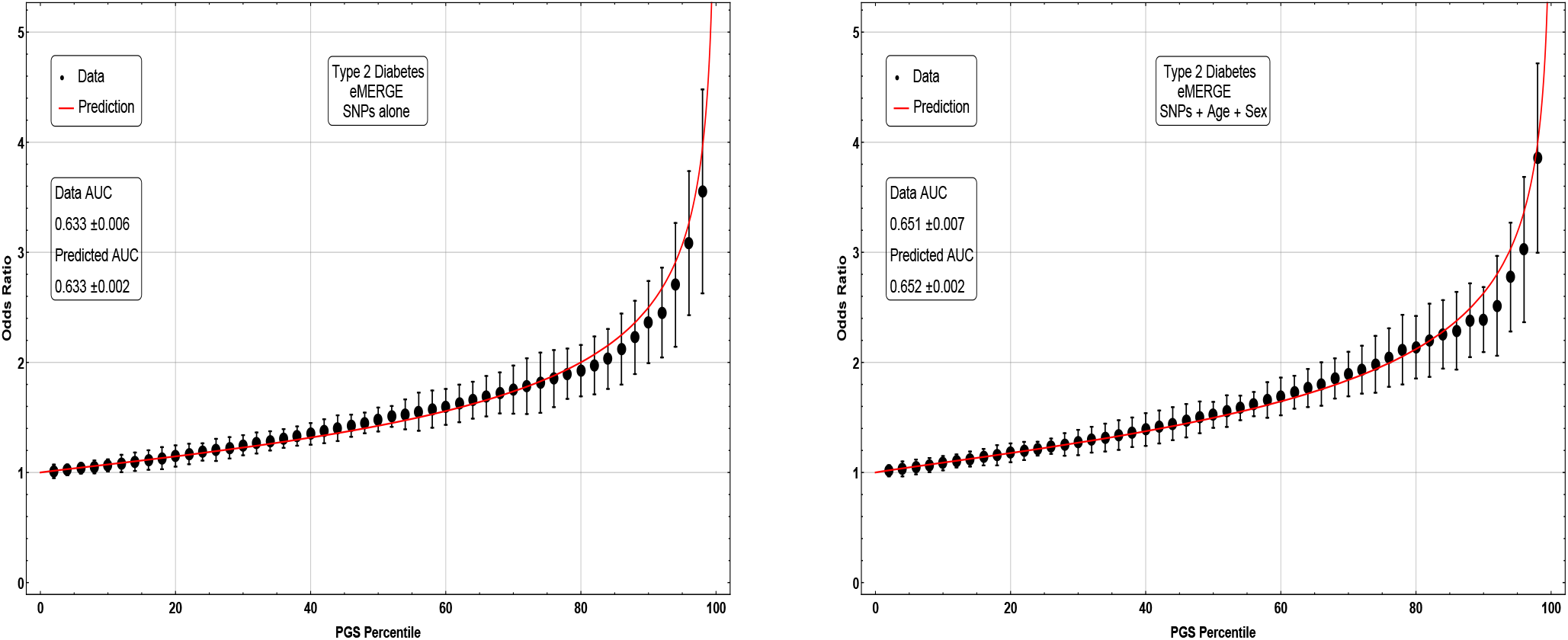
Odds ratio between upper percentile in PGS and total population prevalence in eMERGE for Type 2 Diabetes with and without using age and sex as covariates.

In our analysis we tested whether altering the regressand (phenotype *y*) to some kind of residual based on age and sex could improve the genetic predictor. In all cases we start with *y* = 1,0 for case or control respectively. Then we use the three different regressands:

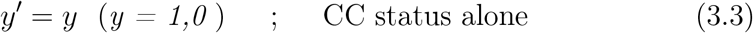

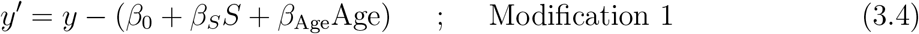

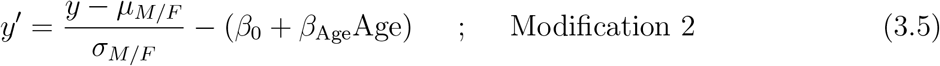

For each case, we tested this including and excluding the sex chromosomes during the regression. As with the previous results, the best prediction accuracy is not appreciably altered if training is done on the autosomes alone. The results are given in table 6.

**Table 6:**
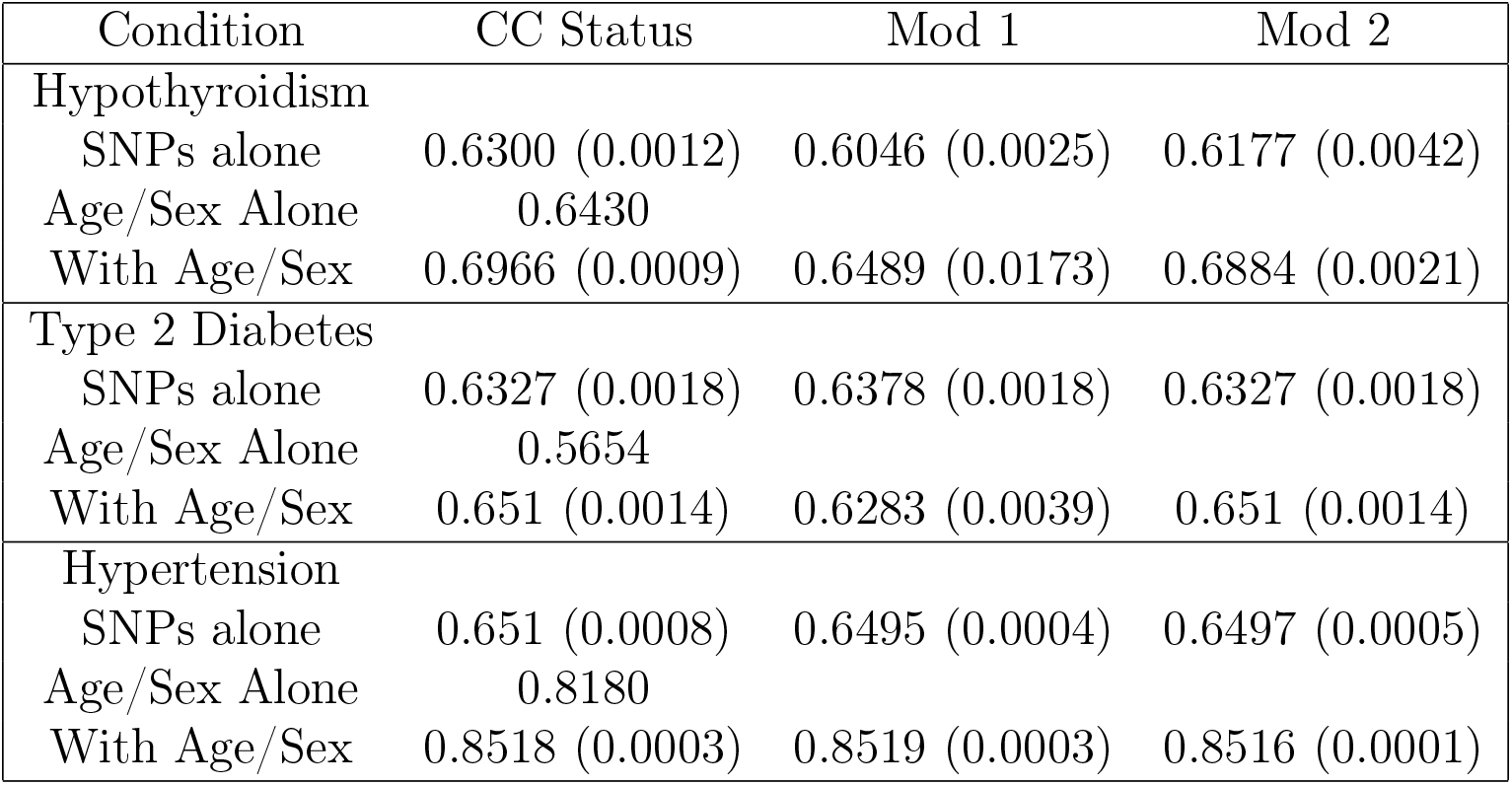
Table of prediction results using three types of regressands. All results are on eMERGE and show results for using SNPs, Age, Sex and combinations of such.

| Condition | CC Status | Mod 1 | Mod 2 |
| --- | --- | --- | --- |
| Hypothyroidism |  |  |  |
| SNPs alone | 0.6300 (0.0012) | 0.6046 (0.0025) | 0.6177 (0.0042) |
| Age/Sex Alone | 0.6430 |  |  |
| With Age/Sex | 0.6966 (0.0009) | 0.6489 (0.0173) | 0.6884 (0.0021) |
| Type 2 Diabetes |  |  |  |
| SNPs alone | 0.6327 (0.0018) | 0.6378 (0.0018) | 0.6327 (0.0018) |
| Age/Sex Alone | 0.5654 |  |  |
| With Age/Sex | 0.651 (0.0014) | 0.6283 (0.0039) | 0.651 (0.0014) |
| Hypertension |  |  |  |
| SNPs alone | 0.651 (0.0008) | 0.6495 (0.0004) | 0.6497 (0.0005) |
| Age/Sex Alone | 0.8180 |  |  |
| With Age/Sex | 0.8518 (0.0003) | 0.8519 (0.0003) | 0.8516 (0.0001) |

The distributions in Figs. 3–5 appear Gaussian under casual inspection, and were further tested against a normal distribution. We illustrate this with Atrial Fibrillation and Testicular cancer – these two conditions represent respectively the best and worst fits to Gaussians. For control groups, results were similar for all phenotypes. For example assuming “Sturge’s Rule” for the number of bins, Atrial Fibrillation controls lead to 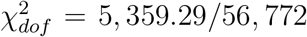 with a p-value 7 × 10^−1013^ when tested against a Gaussian distribution. For cases, we also found extremely good fits. Again, Atrial Fibrillation cases lead to 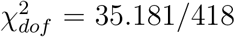 and p-value 0. 0192. Even for phenotypes with very few cases we find very good fits. For Testicular Cancer cases we find a 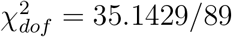 and p-value 1.18 × 10^−4^. For predicted AUCs and Odds Ratios using Eqs. (3.1) & (3.2) we find very little difference between using means and standard deviations from empirical data sets or using fits to Gaussians.

As more data become available for training we expect prediction strength (e.g., AUC) to increase. Based on estimated heritability, predictors in this study are still far from maximum possible AUCs, such as: type 2 diabetes (.94), coronary artery disease (.95), breast cancer (.89), prostate cancer (.90), and asthma (.88) [12]. We investigate improvement with sample size by varying the number of cases used in training. For Type 2 Diabetes and Hypothyroidism, we train predictors with 5 random sets of 1k, 2k, 3k, 4k, 6k, 8k, 10k, 12k, 14k, and 16k cases (all with the same number of controls). For Hypertension, we train predictors using 5 randoms sets of 1k, 10k, 20k,…, and 90k cases. For each, we also include the previously generated best predictors which used all cases except the 1000 held back for crossvalidation. These predictors are then applied to the eMERGE dataset and the maximum AUC is calculated.

In Figure 11 we plot the average maximum AUC among the 5 training sets against the log of the number of cases (in thousands) used in training. Note that in each situation, as the number of cases increases, so does the average AUC. For each disease condition, the AUC increases roughly linearly with log N as we approach the maximum number of cases available. The rate of improvement for Type 2 Diabetes appears to greater than for Hypertension or Hypothyroidism, but in all cases there is no sign of diminishing returns. There is obviously a ceiling to the amount of improvement, determined by the heritability of the specific condition [12], but we see no evidence that we are approaching that limit.

**Figure 11:**
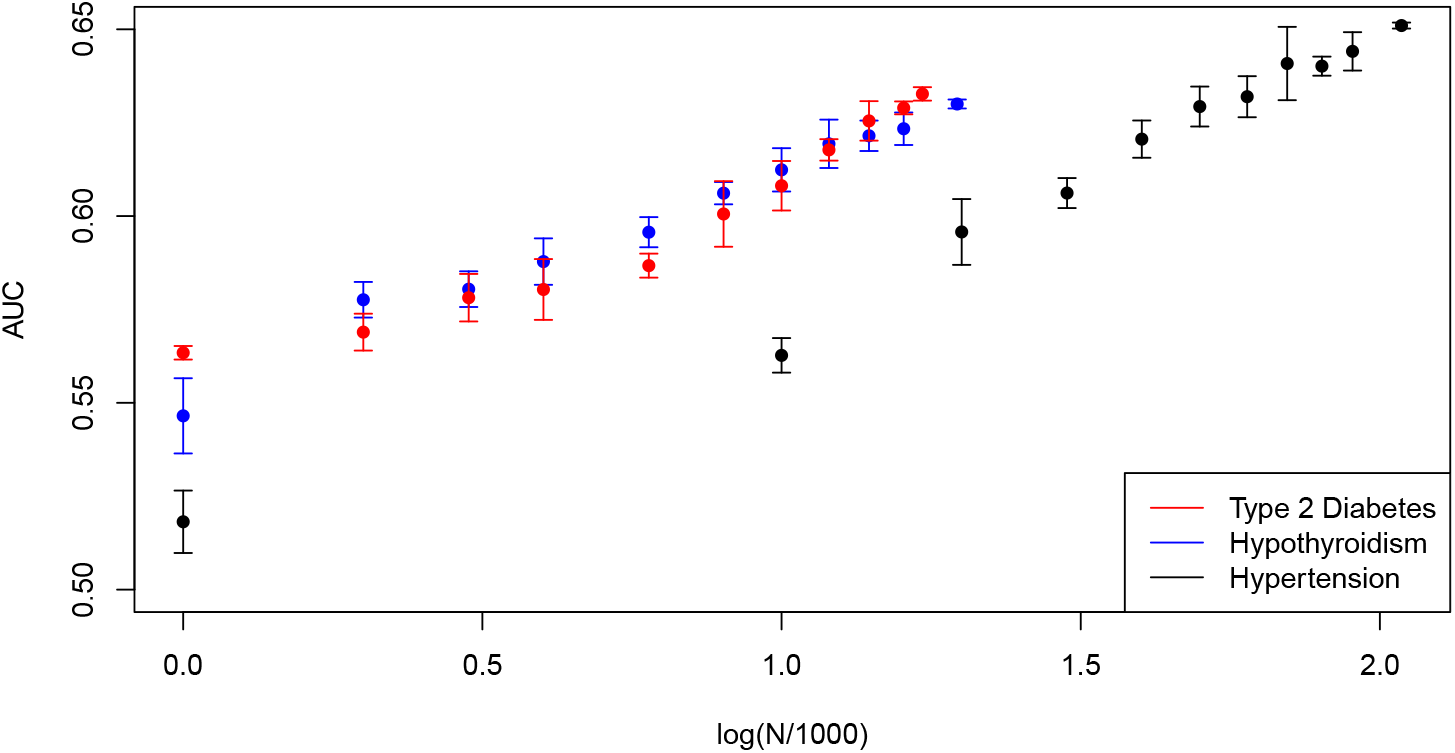
Maximum AUC on out-of-sample testing set (eMERGE) as a function of the number of cases (in thousands) included in training. Shown for type 2 diabetes, Hypothyroidism and Hypertension

By extrapolating this linear trend, we can project the value of AUC obtainable using a future cohort with a larger number of cases. In this work, we trained Type 2 Diabetes, Hypothyroidism and Hypertension predictors using 17k, 20k and 108k cases, respectively. If, for example, cohorts were assembled with 100k, 100k and 500k cases, then the linear extrapolation suggests AUC values of 0.70, 0.67 and 0.71 respectively. This corresponds to 95 percentile odds ratios of approximately 4.65, 3.5, and 5.2. In other words, it is reasonable to project that future predictors will be able to identify the 5 percent of the population with at least 3-5 times higher likelihood for these conditions than the general population. This will likely have important clinical applications, and we suggest that a high priority should be placed on assembling larger case data sets for important disease conditions.

We focused on the three traits above because we can test out of sample using eMERGE. However, using the Ancestry Out of Sample (AOS) method, we can make similar projections for diseases which may 1) be more clinically actionable or 2) show more promise for developing well separated cases and controls. We perform AOS testing while varying the number of cases included in training for Type 1 Diabetes, Gout, and Prostate Cancer. We train predictors using all but 500, 1000, and 1500 cases and fit the maximum AUC to log(*N*/1000) to estimate AUC in hypothetical new datasets. For Type 1 Diabetes, we train with 2234, 1734 and 1234 cases – which achieve AUC of 0.646, 0.643, 0.642. For Gout we train with 5503, 5003 and 4503 cases achieving AUC of 0.0.681, 0.676, 0.0.673. For Prostate Cancer, we train with 2758, 2258, 1758 cases achieving AUC of 0.0.633, 0.628, 0.609. A linear extrapolation to 50k cases of Prostate Cancer, Gout, and Type 1 Diabetes suggests that new predictors could achieve AUCs of 0.79, 0.76 and 0.66 (respectively) based solely on genetics. Such AUCs correspond to odds ratios of and 11, 8, and 3.3 (respectively) for 95th percentile PGS score and above.

## 4 Discussion

The significant heritability of most common disease conditions implies that at least some of the variance in risk is due to genetic effects. With enough training data, modern machine learning techniques enable us to construct polygenic predictors of risk. A learning algorithm with enough examples to train on can eventually identify individuals, based on genotype alone, who are at unusually high risk for the condition. This has obvious clinical applications: scarce resources for prevention and diagnosis can be more efficiently allocated if high risk individuals can be identified while still negative for the disease condition. This identification can occur early in life, or even before birth.

In this paper we used UK Biobank data to construct predictors for a number of conditions. We conducted out of sample testing using eMERGE data (collected from the US population) and Ancestry Out of Sample (AOS) testing using UK ethnic subgroups distinct from the training population. The results suggest that our polygenic scores indeed predict complex disease risk – there is very strong agreement in performance between the training and out of sample testing populations. Furthermore, in both the training and test populations the distribution of PGS is approximately Gaussian, with cases having on average higher scores. We verify that, for all disease conditions studied, a simple model of displaced Gaussian distributions predicts empirically observed odds ratios (i.e., individual risk in test population) as a function of PGS. This is strong evidence that the polygenic score itself, generated for each disease condition using machine learning, is indeed capturing a nontrivial component of genetic risk.

By varying the amount of case data used in training, we estimate the rate of improvement of polygenic predictors with sample size. Plausible extrapolations suggest that sample sizes readily within reach of population genetics studies will result in predictors of significant clinical utility. Additionally, extending this analysis to exome and whole genome data will also improve prediction. The use of genomics in Precision Medicine has a bright future, which is just beginning. We believe there is a strong case for making inexpensive genotyping Standard of Care in health systems across the world.

## Acknowledgments

LL, TR, SY, and SH acknowledge support from the Office of the Vice-President for Research at MSU. This work was supported in part by Michigan State University through computational resources provided by the Institute for Cyber-Enabled Research. The authors are grateful for useful discussion with Steven G. Avery, Gustavo de los Campos and Ana Vasquez. LT acknowledges the additional support of Shenzhen Key Laboratory of Neurogenomics (CXB201108250094A). The authors acknowledge acquisition of datasets via UK Biobank Main Application 15326.

## A Genotype Quality Control

The main dataset used for training in this work is the 2018 release of the UK Biobank (the 2018 version corrected some issues with imputation, included sex chromosomes, etc). In all predictor training, we restricted our analysis to genetically British individuals (as defined using ancestry principal component analysis performed by UK Biobank) [36]. In 2018, the UK Biobank (UKBB) re-released the dataset representing approximately 500,000 individuals genotyped on two Affymetrix platforms – approximately 50,000 samples on the UKB BiLEVE Axiom array and the remainder on the UKB Biobank Axiom array. The genotype information was collected for 488,377 individuals for 805,426 SNPs which were then subsequently imputed to a much larger number of SNPs.

The imputed data set was generated using the set of 805,426 raw markers using the Haplotype Reference Consortium and UK10K haplotype resources. After imputation and initial QC, there were a total of 97,059,328 SNPs and 487,409 individuals. From this imputed data, further quality control was performed using Plink version 1.9. For out-of-sample testing of polygenic risk scores, imputed UK Biobank SNPs which survived the prior quality control measures, and are also present in a second dataset from the Electronic Medical Records and Genomics (eMERGE) study are kept. After keeping SNPs which are common to both the UK Biobank and eMERGE, 557,595 SNPs remained. Additionally SNPs and samples which had missing call rates exceeding 3% were excluded and SNPs with minor allele frequency below 0.1% were also removed so to avoid rare variants. This resulted in 468,514 SNPs and, upon restricting to genetically British, 408,954 people.

## B Phenotype Quality Control

For model training which can be compared to true out-of-sample data, we focused on three case-control conditions which were present in both the UK Biobank and eMERGE datasets – Hypothyroidism, Type 2 Diabetes, and Hypertension. To select Type 2 Diabetes cases in UKBB, we identify individuals based on a doctor’s diagnosis using the fields Diagnoses primary ICD10 or Diagnoses secondary ICD10. Specifically, any individual with ICD10 code E11.0-E11.9 (Non-insulin-dependent diabetes mellitus) in the Main Diagnosis or Secondary Diagnosis field. For training only, we wanted to exclude younger individuals who may still yet develop Type 2 Diabetes, so controls were selected using individuals in the remainder of the UKBB population not identified as cases and born on 1945 or earlier. This resulted in 18,194 cases and 108,726 controls among genetically British individuals.

For both Hypertension and Hypothyroidism, we used the field “Non-Cancer Illness Code (self-reported)” to identify cases and controls. As in the case of type 2 diabetes, we exclude younger individuals as controls for Hypertension. This was not required for Hypothyroidism. Specifically, cases were identified by anyone with the code “1065” (Hypertension) in “Non cancer illness code (self-reported)” and the remainder of the UKBB population who was born before 1950 were selected as controls. This resulted in 109,662 cases and 140,689 controls for Hypertension. For Hypothyroidism, cases were identified by anyone with the code “1226” (Hypothyroidism/Myxoedema) in “Non cancer illness code (self-reported)” and the remainder of the UKBB population was used as a control. This resulted in 20,656 cases and 388,298 controls for Hypothyroidism.

For the following phenotypes we did not have true out of sample data, and so used the Ancestry Out-of-Sample (AOS) based testing procedure of Appendix D: Gout, Testicular Cancer, Gallstones, Breast Cancer, Atrial Fibrillation, Glaucoma, Type 1 Diabetes, High Cholesterol, Asthma, Basal Cell Carcinoma, Malignant Melanoma, Prostate Cancer, and Heart Attack. All conditions were identified using the fields “Non cancer illness code (selfreported)”, “Cancer code (self-reported)” and “Diagnoses primary ICD10” or “Diagnoses secondary ICD10”.

Cases and controls of the following non-cancer illnesses are identified using the field “NonCancer Illness Code (self-reported)”: Gout, Gallstones, Atrial Fibrillation, Glaucoma, High Cholesterol, Asthma and Heart Attack. Cases for a specific non-cancer illness were identified as any individual with the following codes, and the remaining population are selected as a controls: Gout 1466, Gallstones 1162, Atrial Fibrillation 1471, Glaucoma 1277, High Cholesterol 1473, Asthma 1111, Heart Attack 1075. Cases and controls of the following cancer conditions were extracted from the field “Cancer Code (self-reported)”: Testicular Cancer, Prostate Cancer, Breast Cancer, Basal Cell Carcinoma and Malignant Melanoma. Specifically, cases were identified as any individual with the following codes, and controls are the remainder of the population: Testicular Cancer 1045, Breast Cancer 1002, Basal Cell Carcinoma 1061, Malignant Melanoma 1059, Prostate Cancer 1044. To select Type 1 Diabetes cases in UKBB, we identify individuals based on a doctor’s diagnosis using the fields “Diagnoses primary ICD10” or “Diagnoses secondary ICD10”. Specifically, any individual with ICD10 code E10.0-E10.9 (Insulin-dependent diabetes mellitus) in the Main Diagnosis or Secondary Diagnosis field.

After identifying cases and controls in the whole UKBB population, we restricted our training set to “Genetically British” and our testing set to self-reported non-genetically-British whites. The number of cases and controls identified in this manner are listed in Table 7.

**Table 7:** Table of number of cases and controls in training and testing sets for psuedo outof-sample testing. Traits with (*) are trained and tested only on a single sex.

| Condition | Cases (train) | Controls (train) | Cases (test) | Controls (test) |
| --- | --- | --- | --- | --- |
| Gout | 6,003 | 395,842 | 811 | 56,383 |
| Gallstones | 7,022 | 394,823 | 936 | 56,258 |
| Atrial Fibrillation | 3,502 | 398,343 | 420 | 56,774 |
| Glaucoma | 4,609 | 397,236 | 577 | 56,617 |
| Type 1 Diabetes | 2,734 | 399,111 | 388 | 56,806 |
| High Cholesterol | 52,398 | 349,447 | 6,937 | 50,257 |
| Asthma | 47,237 | 354,608 | 6,655 | 50,539 |
| Basal Cell Carcinoma | 4,132 | 397,713 | 577 | 56,617 |
| Malignant Melanoma | 3,301 | 398,544 | 444 | 56,750 |
| Heart Attack | 9,657 | 398,544 | 1,347 | 55,847 |
| Prostate Cancer * | 3,258 | 181,518 | 379 | 24,733 |
| Breast Cancer * | 9,177 | 207,892 | 1,344 | 30,738 |
| Testicular Cancer * | 716 | 184,060 | 91 | 25,021 |

We include a supplementary table which outlines what fraction of cases and controls are male or female. We also include the mean year of birth for male/female cases/controls. This is shown in Table 8.

**Table 8:** Table of fraction of cases and controls and mean year of birth by sex for psuedo out-of-sample testing. Traits with (*) are trained and tested only on a single sex.

| Condition | % Female |  | Mean Birth Year (Female) |  |
| --- | --- | --- | --- | --- |
|  | Cases | Controls | Cases | Controls |
| Gout | 7.35 | 54.98 | 1946.4 | 1951.5 |
| Gallstones | 77.59 | 53.87 | 1949.0 | 1951.6 |
| Atrial Fibrillation | 31.06 | 54.48 | 1945.8 | 1951.5 |
| Glaucoma | 46.91 | 54.36 | 1946.5 | 1951.5 |
| Type 1 Diabetes | 41.45 | 54.36 | 1950.4 | 1951.5 |
| High Cholesterol | 42.98 | 55.95 | 1946.7 | 1952.0 |
| Asthma | 57.48 | 53.85 | 1952.0 | 1951.4 |
| Basal Cell Carcinoma | 58.40 | 54.23 | 1948.5 | 1951.5 |
| Malignant Melanoma | 58.88 | 54.24 | 1949.6 | 1951.5 |
| Heart Attack | 19.68 | 55.11 | 1945.9 | 1951.5 |
| Prostate Cancer * | 0.0 | 0.0 | NA | NA |
| Breast Cancer * | 100.0 | 100.0 | 1946.0 | 1951.6 |
| Testicular Cancer * | 0.0 | 0.0 | NA | NA |
| Condition | % Male |  | Mean Birth Year (Male) |  |
|  | Cases | Controls | Cases | Controls |
| Gout | 92.65 | 45.02 | 1948.5 | 1951.2 |
| Gallstones | 22.41 | 46.13 | 1947.5 | 1951.1 |
| Atrial Fibrillation | 68.94 | 45.52 | 1946.4 | 1951.1 |
| Glaucoma | 53.09 | 45.64 | 1946.5 | 1951.1 |
| Type 1 Diabetes | 58.55 | 45.64 | 1949.0 | 1951.1 |
| High Cholesterol | 57.02 | 44.05 | 1947.3 | 1951.8 |
| Asthma | 42.52 | 46.15 | 1952.0 | 1951.0 |
| Basal Cell Carcinoma | 41.60 | 45.77 | 1947.4 | 1948.5 |
| Malignant Melanoma | 41.12 | 45.76 | 1948.2 | 1951.1 |
| Heart Attack | 80.32 | 44.89 | 1946.2 | 1945.9 |
| Prostate Cancer * | 100.0 | 100.0 | 1944.3 | 1951.2 |
| Breast Cancer * | 0.0 | 0.0 | NA | NA |
| Testicular Cancer * | 100.0 | 100.0 | 1953.2 | 1951.5 |

## C Out-of-sample Quality Control

For out-of-sample testing, we use the 2015 release of the Electronic Medical Records and Genomics (eMERGE) study of approximately 15k individuals available on dbGaP [16]. The specific eMERGE data set used here refers to data downloaded from the dbGaP web site, under accession phs000360.v3.p1.(https://www.ncbi.nlm.nih.gov/projects/gap/cgi-bin/study.cgi?study_id=phs000360.v3.p1). The eMERGE dataset consists of 14,908 individuals with 561,490 SNPs which were genotyped on the Illumina Human 660W platform. The Plink 1.9 software is used for all further quality control. We first filter for SNPs which are common to the UK Biobank. SNPs and samples with missing call rates exceeding 3% are excluded and SNPs with minor allele frequency below 0.1% were also removed. This results in 557,595 SNPs and 14,906 individuals. Of these, the 468,514 SNPs which passed QC on the UK Biobank are used in training.

All eMERGE individuals in our dataset have self reported their ethnicity as white. Not all individuals in eMERGE are strictly cases or controls for any one particular condition. For Type 2 Diabetes, there are 1,921 identified cases and 4,369 identified controls. For Hypothyroidism there are 1084 identified cases and 3171 identified controls. For Hypertension, as the study focused on identifying individuals with Resistant Hypertension, there are two types of cases and two types of controls. Case group 1 consists of subjects with 4 or more medications simultaneous on at least 2 occasions greater than one month apart. Case group 2 has two outpatient (if possible) measurements of systolic blood pressure over 140 or diastolic blood pressure greater than 90 at least one month after meeting medication criteria while still on 3 simultaneous classes of medication AND has three simultaneous medications on at least two occasions greater than one month apart. Control group 2 consists of subjects with no evidence of Hypertension. Control group 1 consists of subjects with outpatient measurements of SBP over 140 or DBP over 90 prior to beginning medication AND has only one medication AND has SBP < 135 and DBP < 90 one month after beginning medication. For model testing of Hypertension, we classified case group 1, case group 2 and control group 1 as cases, while control group 2 is used as controls. For Resistant Hypertension, we classified case group 1 and case group 2 as cases, while control group 2 is used as controls – control group 1 is excluded from this testing. The size of the self-reported white members of the groups are: case group 1 – 952, case group 2 – 406, control group 1 – 677, control group 2 – 1202.

The year of birth in eMERGE is given by decade, so the year of birth is taken to be the 5th year of the decade (i.e., if the decade of birth is 1940, then 1945 is used as year of birth). Some individuals did not have a year of birth listed – these individuals are included when testing models which did not feature age and sex as covariates, but are excluded when testing a model which included age. For obtaining age and sex effects, we used the entire UK Biobank for training as opposed to excluding younger participants as was done for the genetic models.

## D Testing using Genetically Dissimilar Subgroups: Ancestry Out-Of-Sample Testing

For many case-control phenotypes, we do not have access to a second data set for proper out-of-sample testing. For these traits, we follow an ancestry out-of-sample (AOS) testing procedure which was proposed and used in [17]. In this procedure, the predictor is trained on individuals of a homogeneous ethnic background: from UKBB we use genetically British individuals, defined using principal components analysis of population data. The predictor is then applied to individuals who are genetically dissimilar to the training set but not overly distant. For our testing set we use self-reported white (i.e., European) individuals (British/Irish/Any Other White) who are not in the cohort identified as genetically British. These individuals might be, for example, people of primarily Italian, Spanish, French, German, Russian, or mixed European ancestry who now live in the UK.

To identify the genetically British individuals, we follow the procedure in [36]. The top 20 principal components for the entire sampled population are provided directly from UK Biobank and the top 6 are used to identify genetically British individuals. We select individuals who self-report their ethnicity as “British” and use the outlier detection algorithm from the R-package “Aberrant” [55] to identify individuals using pairs of principal component vectors.

Aberrant uses a parameter which is the ratio of standard deviations of the outlying to normal individuals (λ) (Note λ here is a variable name used in Aberrant. It should not be confused with the lasso penalization parameter used in our optimization). This parameter is tuned to make a training set which is overly homogenous compared to those reported as genetically British by the UKBB (λ ~ 20). Because Aberrant uses two inputs at a time, individuals to be excluded from training were identified using principal component pairs (first and second, third and fourth, fifth and sixth) and the union of these sets are the total group which is excluded in the final training set. There were a total of 402,937 individuals to be used in training after principal component filtering.

For this type of testing, the directly called gentoypes are used for training, cross-validation and testing (imputed SNPs are only used for true out-of-sample testing). First, only selfreported white individuals were selected (472,856) and then SNPs and samples with missing call rates exceeding 3% were removed, as were SNPs with minor allele frequency below 0.1% (all using Plink). This results in 658,543 SNPs and 459,039 total individuals which consists of 401,845 genetically British who are used for training and 57,194 non-British self-reported white individuals are used for final ancestry based out-of-sample testing.

### Odds Ratios for AOS

We collect here the odds ratio cumulant plots as a function of PGS percentile (i.e., a given value on the horizontal axis represents individuals with that PGS *or higher*) for the various phenotypes that were tested with the AOS procedure described in Sec. 2 and reported in Table 1. We also comment here on some of the more notable comparisons to previous methods used in the literature to analyze the genetic predictability of these phenotypes. It should be noted that some of these phenotypes – e.g., Asthma, Heart Attack, and High Cholesterol – have been heavily linked to other complex traits and future studies using multiple complex traits might greatly improve prediction.

Asthma, in Fig. 12, has long been known to have a significant genetic component. In this study odds ratios ~ 3x are found for people with PGS scores in the 96^th^ percentile and above. This compares favorably to the literature where 2.5x odds ratio increase at 95% confidence level was found for children with parents that have asthma [56]. GWAS studies [57] have shown that Asthma seems to be correlated with both hay fever and Eczema conditions. Although in performing this study, we did not find a strong predictor for Eczema, relevant data is available in UKBB and multi-phenotype studies could be performed in the future.

**Figure 12:**
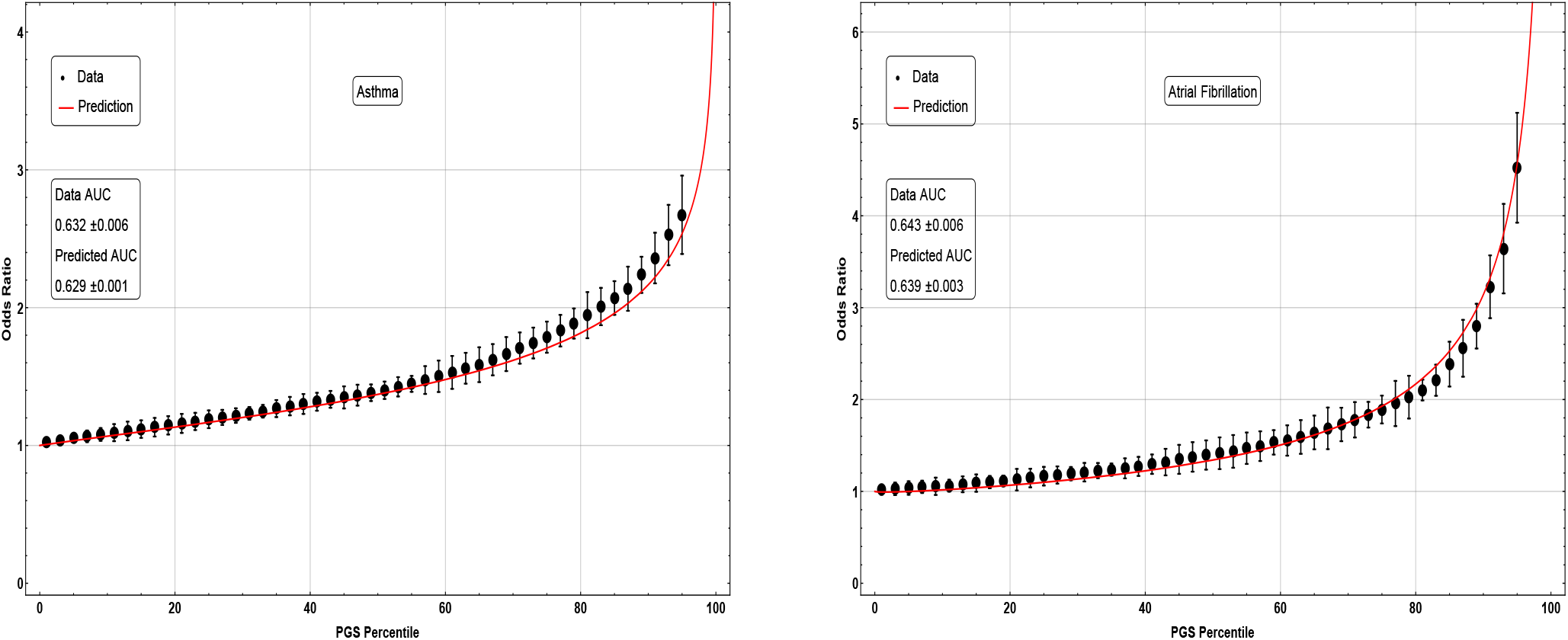
Odds ratio as a function of PGS percentile (i.e. scores at that point or above) for (left) Asthma and (right) Atrial Fibrillation.

Atrial Fibrillation, seen in Fig. 12, is also known to have a genetic risk factor. Parental studies have shown a 1.4x odds ratio, but-although gene loci have been identified, genetic studies have not previously been successful in clinical settings [58]. In this work, PGS scores in the 96^th^ percentile and above predict up to a 5x increase in odds.

Breast Cancer, in Fig. 13, has long been evaluated with the understanding that there is a genetic risk component. Recent studies involving multi SNP prediction (77 SNPs) have been able to predict 3x odds increases for genetic outliers. This is consistent with our results for the highest genetic outliers although we used many more SNPS 480 ± 62.

**Figure 13:**
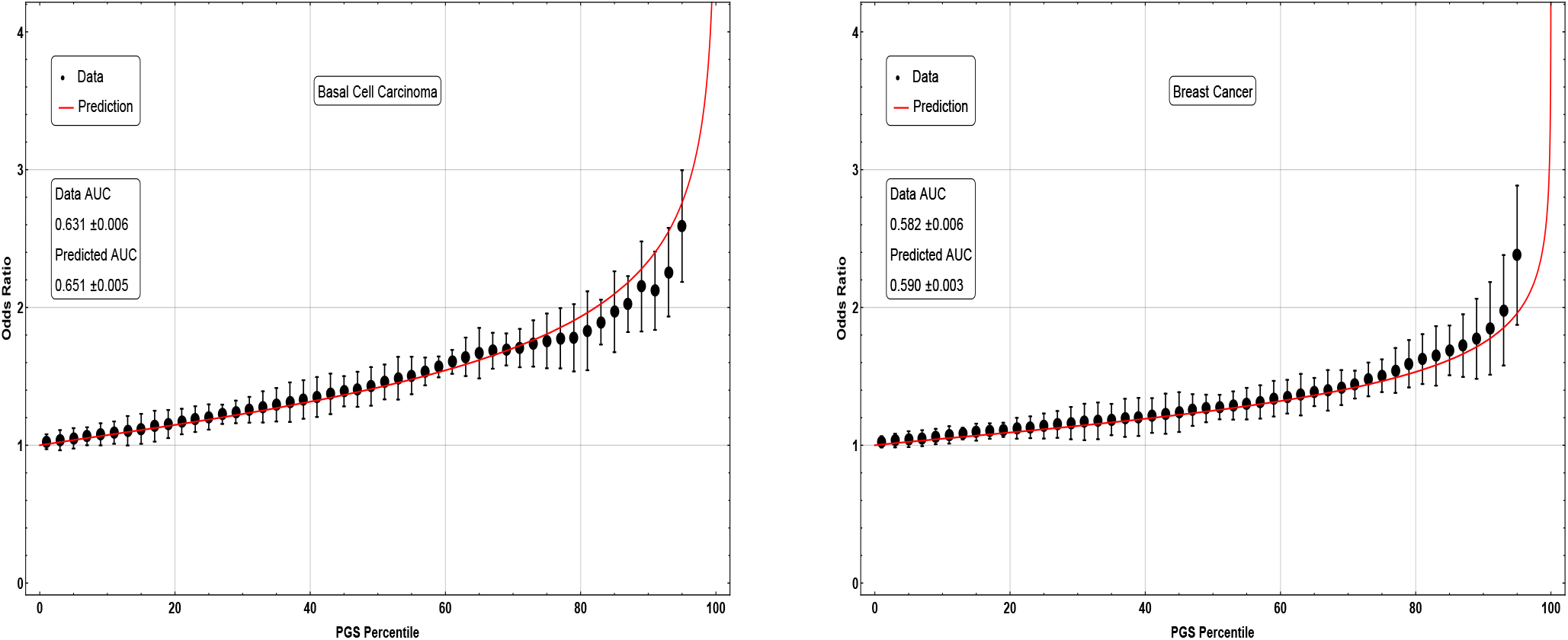
Odds ratio as a function of PGS percentile (i.e. scores at that point or above) for (left) Basal Cell Carcinoma and (right) Breast Cancer.

Recent reviews suggest that much of the risk leading to a higher probability of having Gallstones is associated with non-genetic factors. However, in Fig. 14, we find that 90^th^ percentile and above PGS is be associated with a 3x odds increase.

**Figure 14:**
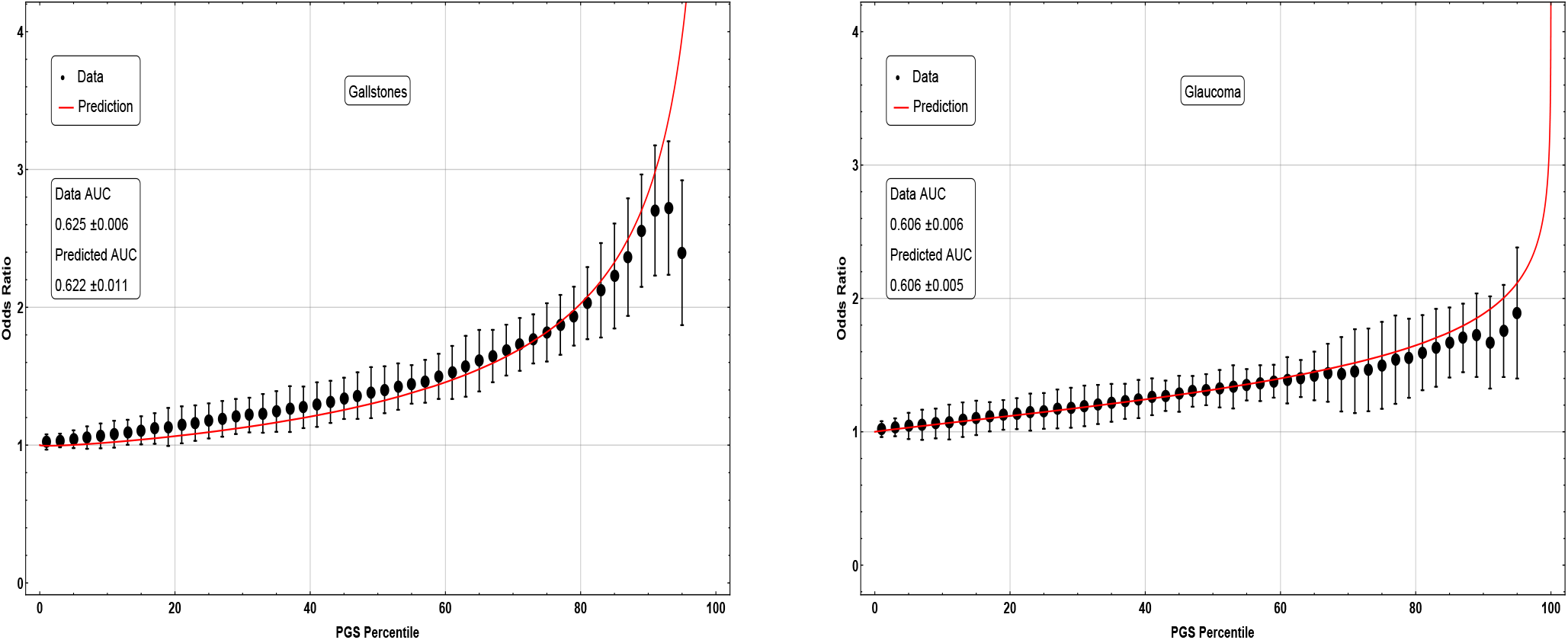
Odds ratio as a function of PGS percentile (i.e. scores at that point or above) for (left) Gallstones and (right) Glaucoma.

While there are a variety of relevant environmental factors, recent reviews of the genetics of Glaucoma [59] highlight that GWAS studies have found 25 genic regions with odds ratios above 1x. The highest being 2.80x [60]. In Fig. 14 we see similar odds ratios for extreme PGS.

Gout, seen in Fig. 15, has an extremely high 4.5x odds ratio for PGS in the *96^th^* percentile and above. Reviews of Gout [61] have noted both a strong familial heritability and known GWAS loci, but we are not aware of previously-computed odds ratios this large solely due to genetics.

**Figure 15:**
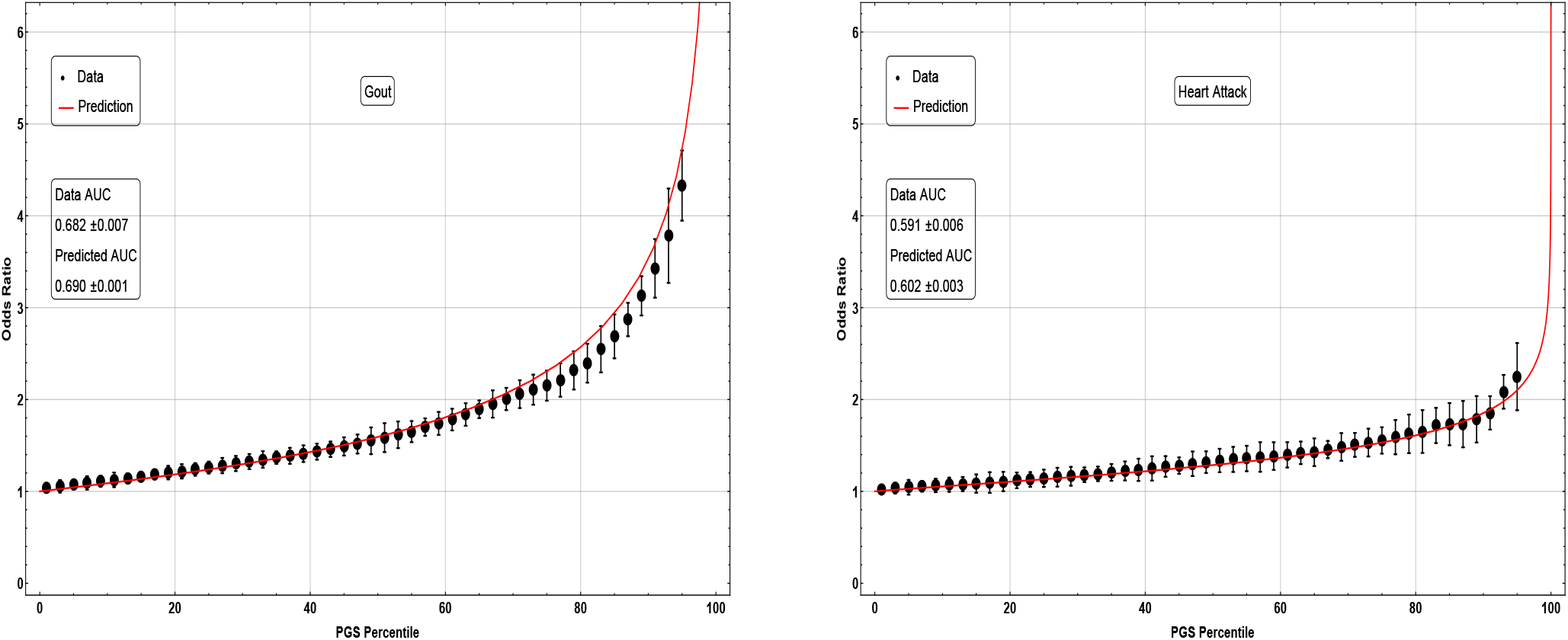
Odds ratio as a function of PGS percentile (i.e. scores at that point or above) for (left) Gout and (right) Heart Attack.

There is a wide ranging literature covering the genetics and heritability of Type 1 Diabetes. In Fig. 17 we see a large 4.5x odds ratio for extreme PGS. Notably, the literature has identified genetic prediction to be extremely useful in differentiating between Type 1 and Type 2 Diabetes [62] and in identifying *β* cell autoimmunity [63], which is highly correlated with diabetes.

**Figure 16:**
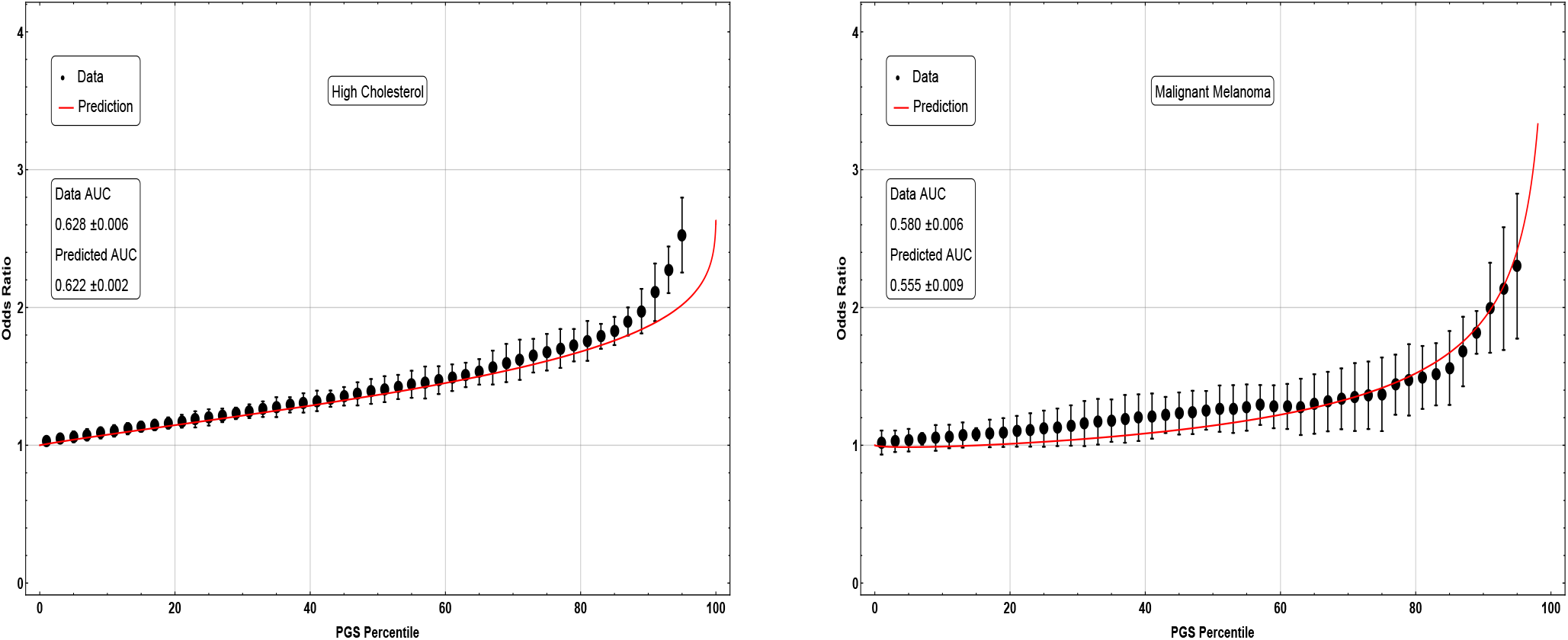
Odds ratio as a function of PGS percentile (i.e. scores at that point or above) for (left) High Cholesterol and (right) Malignant Melanoma.

**Figure 17:**
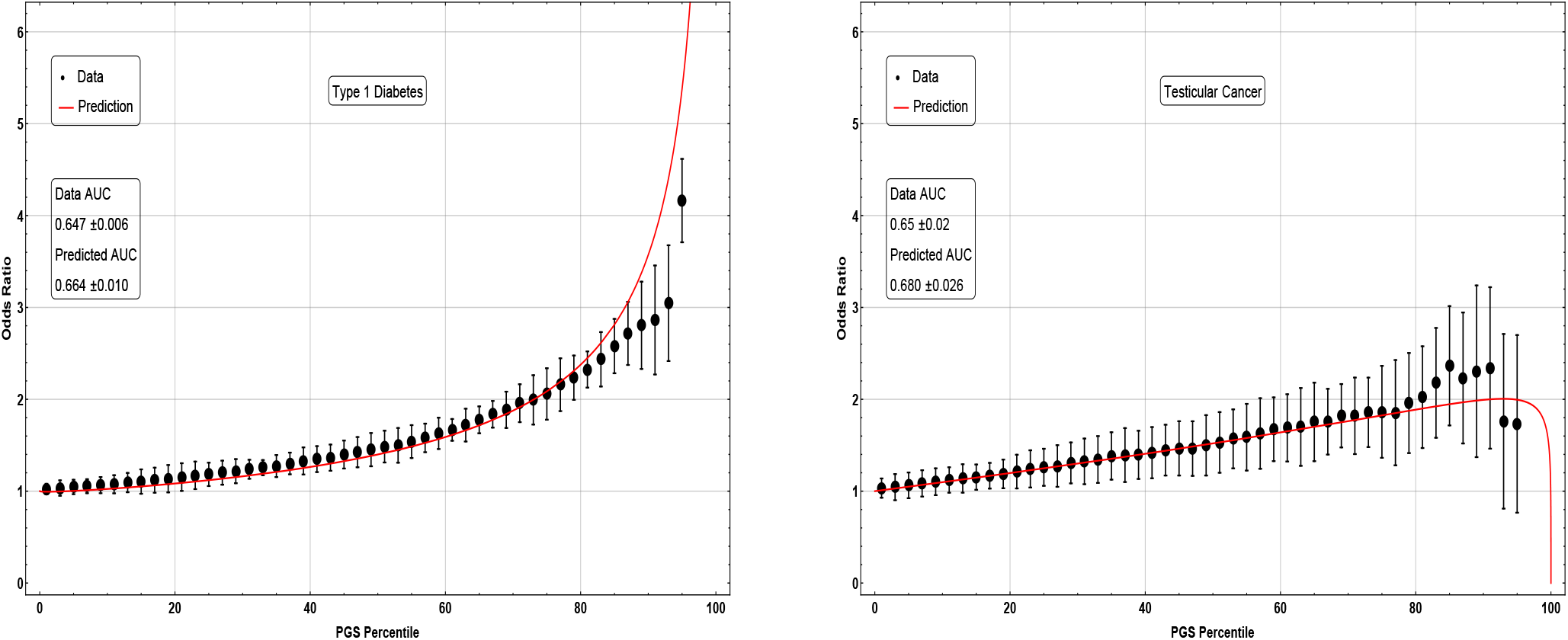
Odds ratio as a function of PGS percentile (i.e. scores at that point or above) for (left) Type 1 Diabetes and (right) Testicular Cancer. Note that the dip at extreme PGS values in the predicted Testicular Cancer curve may be related to a small number of available cases; the cases and controls are not well fit by two separate Gaussian distributions.

Prostate Cancer is the most common gender specific cancer in men. The odds ratio for AOS testing can be seen in Fig. 18. It has long been known that age is a significant risk factor for prostate cancer, but GWAS studies have shown that there is a significant genetic component [64], Additionally, it has been shown, using genome wide complex trait analysis (GCTA), that variants with minor allele frequency 0.1 — 1% make up an important contribution to “missing heritablity” for men of African ancestry [65]. This study includes some SNP variants with minor allele frequency as low as 0.1%, so our model might include some of this contribution.

**Figure 18:**
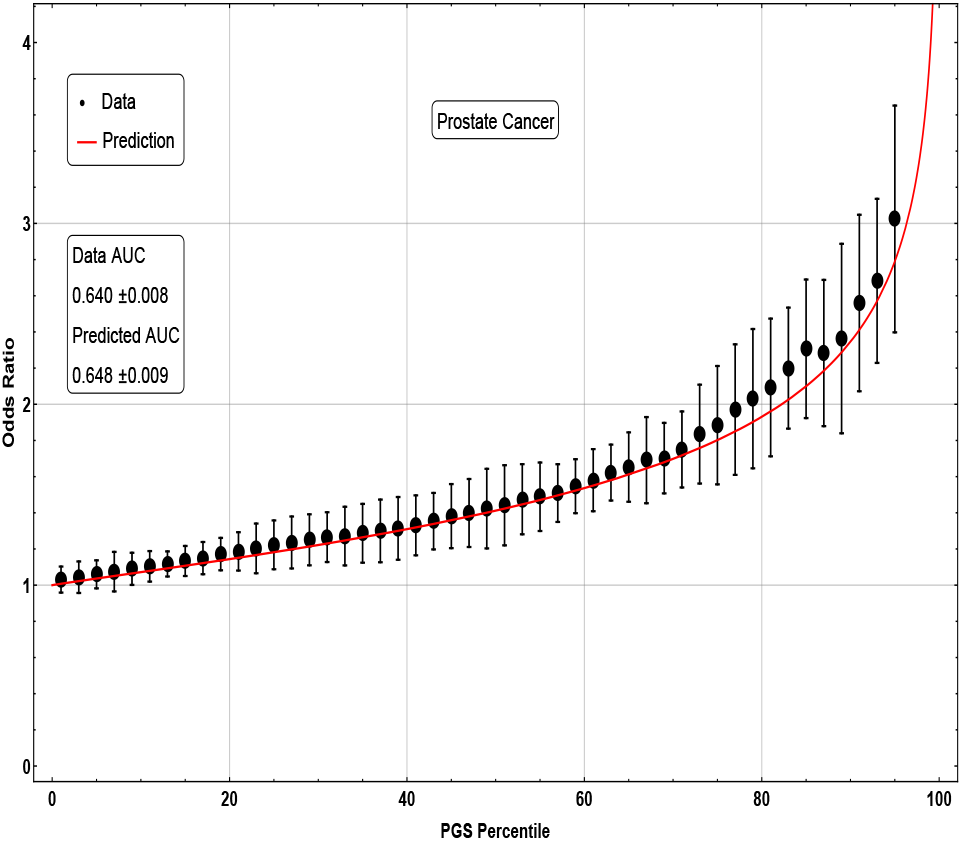
Odds ratio as a function of PGS percentile (i.e. scores at that point or above) for Prostate Cancer.

## E Model Training Algorithm

In all calculations, we use a custom implementation of LASSO regression (Least Absolute Shrinkage and Selection Operator) written in the Julia language. This is the same implementation used in [9]. Given a set of samples *i* = 1, 2,…, *n* with a set of *p* SNPs, the phenotype *y_i_* and state of the *j*^th^ SNP, *X_ij_*, are observed. *X_ij_* is an *n* x *p* matrix which contains the number of copies of the minor allele and any missing values are replaced with the SNP average. The *L*_1_ penalized regression, LASSO, seeks to minimize the objective function

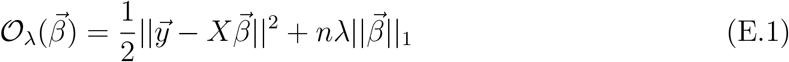

where 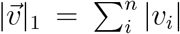 is the *L*_1_ norm, 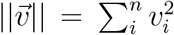 is the *L*_2_ norm and λ is a tuneable hyperparameter. The solution is given in terms of the soft-thresholding function as

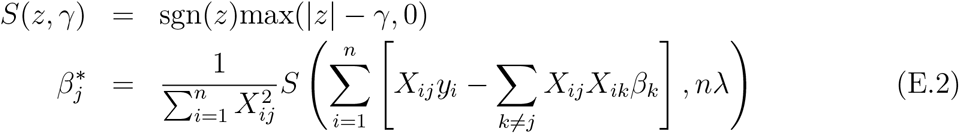

The penalty term affects which elements of 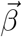 have non-zero entries. The value of λ is first chosen to be the maximum value such that all *β_i_* are zero, and it is then decreased, allowing more nonzero components in the predictor. For each value of 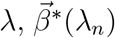 is obtained using the previous values of 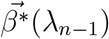 (warm start) and coordinate descent. The Donoho-Tanner phase transition [46] describes how much data is required to recover the true nonzero components of the linear model and suggests that we expect to recover the true signal with s SNPs when the number of samples is n ~ 30s – 100s (see [45, 50]). For a more complete description of the algorithm, see [9].

For all three conditions which are available in eMERGE, we withhold a subset of 1000 cases and 1000 controls from the training set to be set aside for cross-validation. We repeated this 5 times with non-overlapping cross-validation sets. With training and cross-validation sets constructed, we first perform a GWAS on the training set and select the rank ordered top 50,000 p-value SNPs. We then use these SNPs as input to the LASSO algorithm and finally apply the predictor to the corresponding cross-validation set in order to select the value of λ. For conditions which AOS testing is used, we use cross-validation sets of 500 cases and 500 controls to tune our model.

Because individual SNPs are uncorrelated to year of birth and sex, we are able to regress on SNPs independently of age and sex. To train combined models, which include SNPs, age and sex, we perform LASSO on SNPs alone and least squares regression on age and sex only, then add the two predictor scores together. We tested for whether an improvement in AUC is achieved through a simultaneous regression using polygenic score (PGS), age, and sex as covariates, but found this to give similar AUC as doing the regressions independently and adding the results (to within a few % accuracy).

## F Analytic AUC and Risk

While much of this section is well understood, we include a summary to define terminology and for reference. By assuming that cases and controls have PGS distribution which is Gaussian, we can analytically calculate quantities for genetic prediction. For example, we can calculate an AUC and see how it corresponds to to an odds ratio for various distributional parameters. For additional discussion, we refer the interested reader to [54].

Assume a case-control phenotype and that the cases and controls have Gaussian distributed PGS. Letting *i* = {0,1} represent controls and cases respectively, the distribution of scores can be written

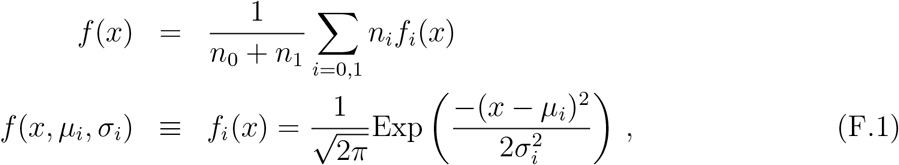

and *n_i_* represents the total number of cases/controls. For completeness, we recall the defintion of the error function here

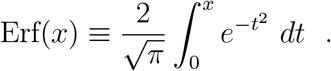

### AUC

First we need to generate an ROC curve of *false positive rate* (FPR) vs *true positive rate* (TPR).

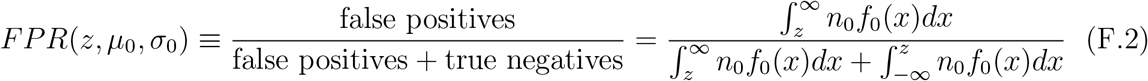

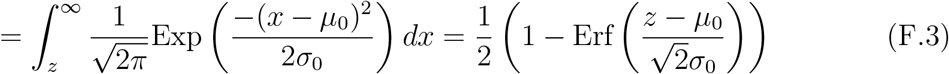

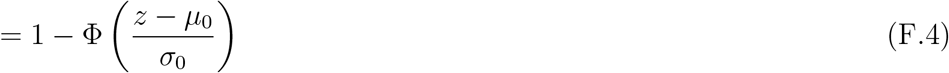

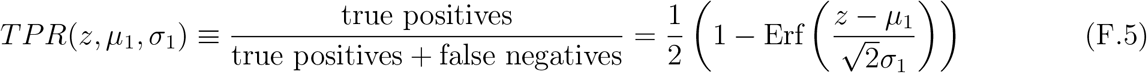

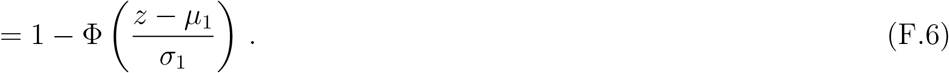

The AUC is then defined as the area under the ROC curve,

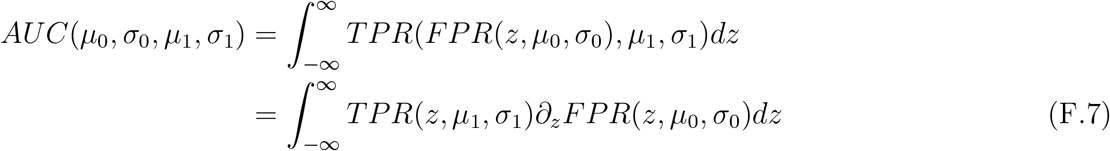

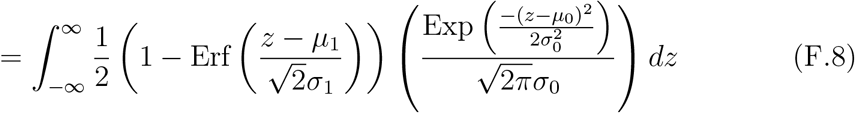

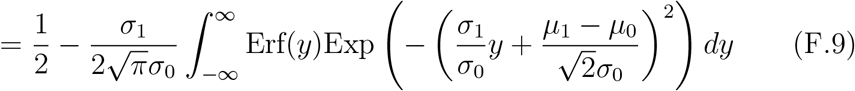

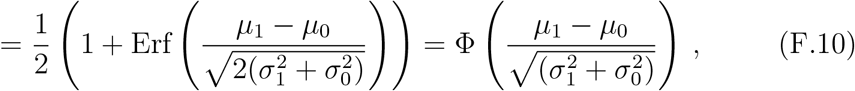

in agreement with Eq.(3.1). Note that the AUC is independent of the number of cases and controls.

### Risk and Odds

There are two standard ways in the literature to classify the increased likelihood of a disease at a higher z-score.

#### Risk Ratio

represents the ratio between (a) the number of cases at a particular z-score and above over the total number of people at z-score and above to (b) the total number of cases over the total number of cases and controls.

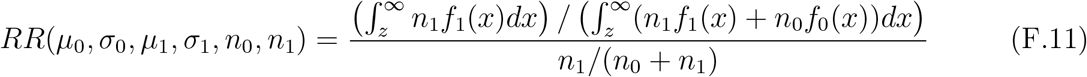

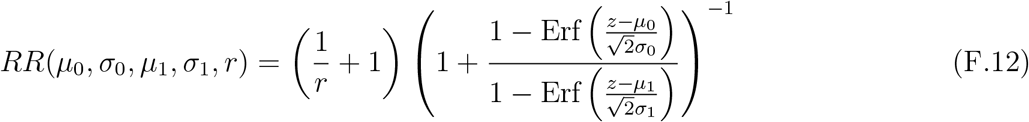

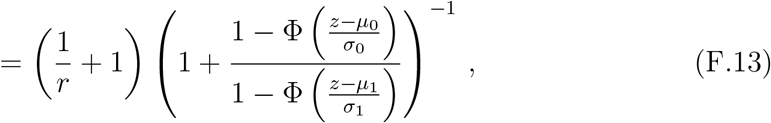

where we can note that the Risk Ratio only depends on the ratio *r* ≡ *n*_1_/*n*_0_.

#### Odds Ratio

represents the ratio between (a) the number of cases at a particular z-score and above over the number of controls at a particular z-score and above to (b) the total number of cases over the total number of controls

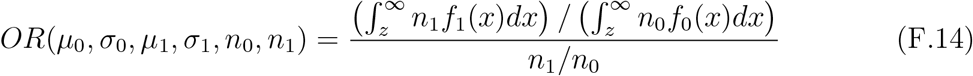

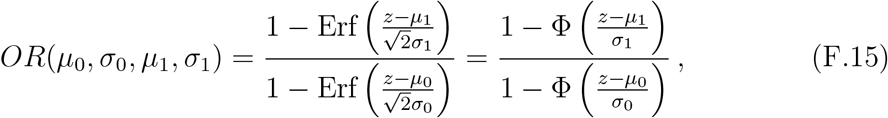

which is *independent* of *n*_1_ and *n*_0_. This is the result Eq.(3.2). Note that in the *rare disease limit* (RDL)

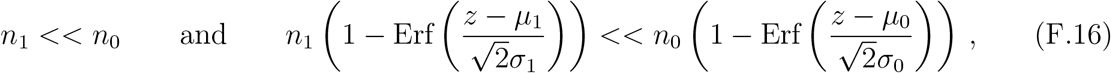

the risk ratio and odds ratio agree

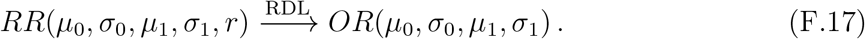

#### PGS percentile

In either case, we would like to know the risk or odds ratio in terms of the percentage of people with a particular z-score and above. We can define this percentile function as

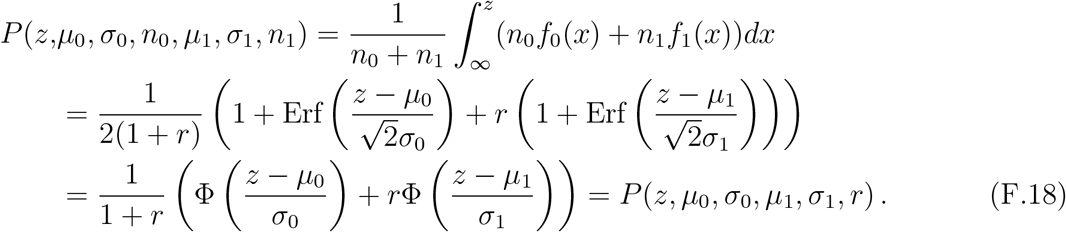

Combining Eq.(3.1), Eq.(3.2), and Eq.(F.18) we can plot the odds ratio in terms of the distributional parameters as seen in Fig. 19.

**Figure 19:**
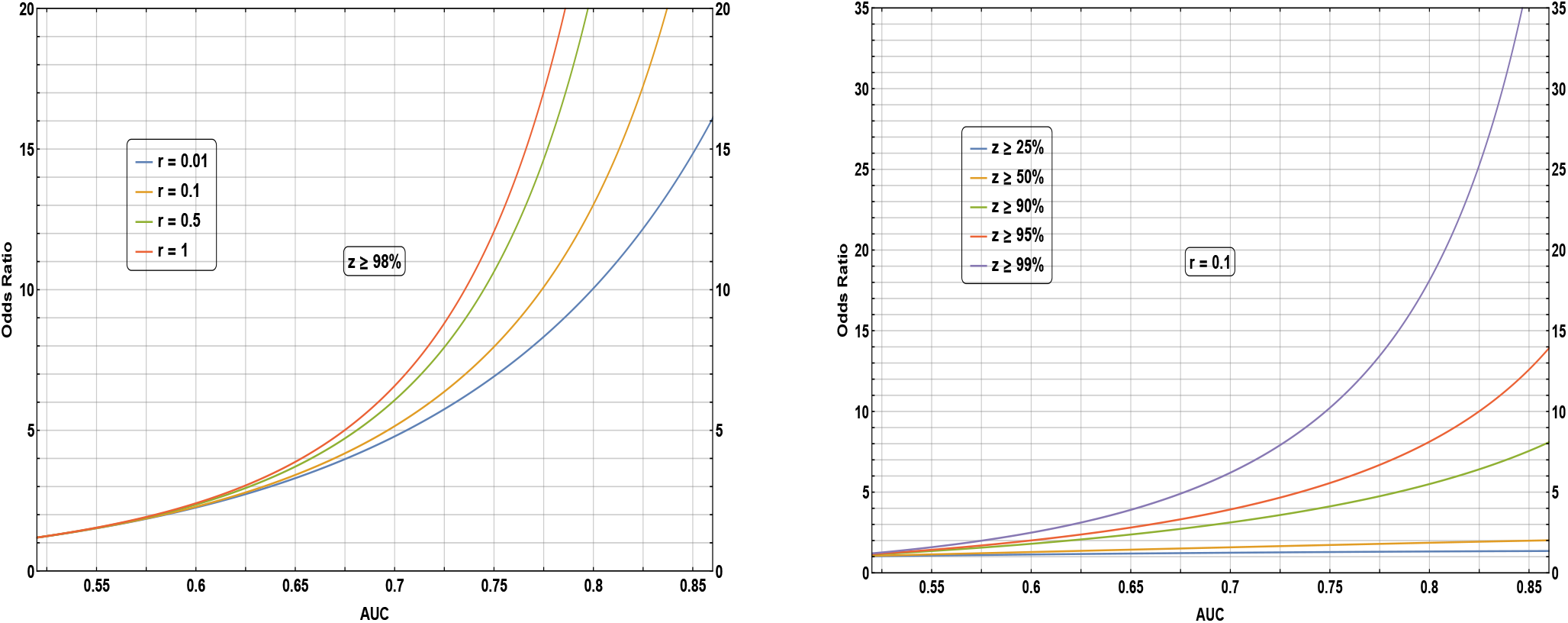
Odds ratio (assuming two displaced Gaussian distributions) as a function of AUC. Left: for z-scores above the 98^th^ percentile at various values of the ratio of cases to controls r. Right: for case to control ratio r = 0.1 at various z-score percentiles. Assuming a population-representative sample, r is the prevalence of the disease in the general population.

## Footnotes

1 The promise of genetic prediction of human complex traits has been discussed for years [3–8], but the use of genome wide predictors for common phenotypes has yet to become commonplace.

2 The 2018 version corrected some issues with imputation, included sex chromosomes, etc. See the Appendices A and B for further details.

3 The AOS testing is a procedure similar to that described in [17] and *not* a detailed analysis of predictor power fall off as a function of ancestry genetic distance. We intend to report on such effects in a future study.

4 There has been some attention to *non-linear* models for complex trait interaction in the literature [37–40]. However we limit ourselves here to additive effects, which have been shown to account for most of the common SNP heritability for human phenotypes such as height [9], and in plant and animal phenotypes. [41–43]

5 In this context, a more parsimonious model refers to one with fewer active SNPs.

6 This is the given accuracy *at a specific number of cases and controls*. As described in Sec. 3 the absolute value of AUC depends on the number of reported cases.

7 The details of the following calculations are in Appendix F. Some of the results can be found in [54].

